# Mechanisms regulating combination effect of antibody-drug conjugates and cancer immunotherapy

**DOI:** 10.64898/2026.07.17.738956

**Authors:** Sehmus Tohumeken, Ali Mostafa, Rashmi Binjawadagi, Mimi Mai, Ana Paucarmayta, Ana Maria Mansilla Merlano, Chistine Youn, Eric Chang, Pooja Shah, Huiyanangel Chow, Wileashire Moulton, Xiaokang Luo, Kerlene Berwick Tam, Matthew Flynn, Leslie Wetzel, Even Walseng, Elizabeth Galery, Joseph Boland, Anna Huntley, Christine Kiefer, Jason Zhang, Carolina Mendoza-Topaz, Corinne Cayatte, Cristina Bergamaschi, Bilal Omar, Puja Sapra, Mark Cobbold, Emilio Sanseviero, Dmitry Gabrilovich

**Affiliations:** Oncology R&D, ICC AstraZeneca Gaithersburg, USA; Biologics Engineering, R&D, AstraZeneca, Gaithersburg, MD, USA; Discovery Sciences, BioPharmaceuticals R&D, AstraZeneca, Gaithersburg, MD, USA; Oncology R&D, OTD AstraZeneca Gaithersburg, USA; Oncology R&D, TM, AstraZeneca, Gaithersburg, MD, USA; Discovery Sciences, BioPharmaceuticals R&D, AstraZeneca Cambridge, UK

**Author notes:** Co-senior Authors. **Conflict of interest** All authors are employees and shareholders of AstraZeneca.

## Abstract

Antibody–drug conjugates (ADCs) have emerged as a transformative class of cancer therapeutics with important challenges still to be addressed. Combination of ADC with immunotherapy is a promising strategy but mechanisms and effective application remain to be determined. We evaluated ADC combinations with T cell engagers (TCEs) and checkpoint inhibitors (CPI). ADC–TCE combinations produced robust antitumor activity independent of antigen and payload and persisted despite ADC-related T cell loss. Efficacy was dominated by a direct effect of ADC on tumor cells. ADCs induced autophagy that upregulated TNF receptors (TNFRs) and mannose-6-phosphate receptors (M6PR). When ADCs were combined with TCEs TNFα released by T cells was primarily responsible for potent antitumor effect of combination. In contrast, M6PR was dispensable for ADC–TCE activity but critical for combinations with CPI expanded antigen-specific T cells via enhanced granzyme B uptake. These data reveal a unifying, target- and payload-agnostic mechanism enabling rational ADC–immunotherapy combinations.

**Significance:** This is first evidence that ADC-induced tumor cell autophagy via up-regulation of TNFR and M6PR could be responsible for potent antitumor effect of combination of ADC with TCE. TCEs exploit a TNFα–TNFR axis, whereas antigen-specific T cells leverage granzyme B–M6PR uptake. This mechanistic framework explains broad ADC–TCE synergy and guides rational selection of ADC–immunotherapy combinations beyond checkpoint blockade.

## Introduction

Over the past decade, antibody-drug conjugates (ADCs) have emerged as a transformative class of cancer therapeutics, combining the specificity of monoclonal antibodies with the cytotoxic potency of chemotherapeutic payloads to deliver targeted cell killing while minimizing systemic toxicity (1). HER2 targeting ADC Enhertu and ADCs targeting trophoblast cell surface antigen 2 (TROP2), a transmembrane glycoprotein overexpressed in various epithelial cancers have been approved by FDA. Datopotamab deruxtecan, TROP2-targeting ADC utilizing a topoisomerase I inhibitor (Top-1i) payload has demonstrated potent antitumor activity with ongoing clinical trials in non-small cell lung cancer (NSCLC) and other solid tumors (2). Sacituzumab govitecan, a TROP2-directed ADC conjugated with payload, has received FDA approval for metastatic triple-negative breast cancer and urothelial carcinoma. Other payloads, such as monomethyl auristatin E (MMAE), a microtubule-disrupting agent, have also shown efficacy in ADC platforms, exemplified by brentuximab vedotin and enfortumab vedotin, which target CD30 and Nectin-4, respectively (3,4). Despite these remarkable advances there are still substantial challenges. ADC efficacy can be constrained by heterogeneous target antigen expression, limited tumor penetration, development of drug resistance, and dose-limiting toxicities that prevent escalation to fully effective doses. Therefore, combining ADC with orthogonal modalities is a promising approach to enhance the effect of therapy.

T-cell engagers (TCE) have emerged as a powerful immunotherapeutic modality capable of redirecting cytotoxic T cells to tumor cells. These bispecific antibodies typically contain one arm targeting CD3 on T cells and another targeting a tumor-associated antigen, creating an immunological synapse that triggers T-cell activation and tumor cell lysis (5). Checkpoint inhibitors targeting the PD-1/PD-L1 axis have become standard-of-care in multiple tumor types, enhancing T-cell effector function by blocking inhibitory signaling pathways. Immunotherapies, however, while capable of inducing durable responses in some patients, are hampered by limited response rates (typically 15-30% in unselected populations), primary and acquired resistance mechanisms, immunosuppressive tumor microenvironments, and immune-related adverse events. ADC as powerful tool to debulk tumor and immunotherapies able to mobilize adaptive immune responses coupled together with the possibility of targeting several tumor antigens simultaneously, has prompted investigation of rational combination strategies. Combination of ADC with check-point inhibitors demonstrated clinical activity. Enfotumab vedotin - Nectin-4 targeting ADC in combination with PD-1 antibody pembrolizumab demonstrated strong clinical activity in muscle-invasive bladder cancer(6) and head and neck cancer(7). Clinical data matched pre-clinical observation (8–10). Combination of ADC with TCE were proposed (11,12) and recent data report first promising result of combination of CD79b targeting ADC (Polatuzumab Vedotin) with TCE (mosunetuzumab or glofitamab) in patients with lymphoma (13,14). In solid tumors the data are lacking.

Despite promise, the concerns exist regarding potential antagonism, particularly if cytotoxic payloads from ADCs inadvertently kill activated T cells within the tumor microenvironment, potentially negating the benefits of immunotherapy. The precise molecular mechanisms underlying their interaction whether synergistic, additive, or antagonistic, remain poorly understood, and the optimal strategies for combining these agents have not been elucidated. Critical questions persist regarding optimal dosing sequences, the impact of different ADC payloads on immune cell function, and whether specific tumor or immune microenvironment features predict response to combination therapy. Furthermore, the molecular and cellular events occurring at the tumor-immune interface following combination treatment have not been characterized.

In this study, we sought to determine whether ADC-TCE combinations produce synergistic antitumor activity; elucidate the molecular mechanisms underlying any observed combination effects; characterize the impact of ADC treatment on tumor-infiltrating T cells; identify key mediators of combination efficacy; and assess whether findings are generalizable across different ADC payloads, target antigens, and forms of T-cell-directed immunotherapy. We identified a mechanism by which ADCs potentiate T-cell-mediated tumor killing through autophagy-dependent upregulation of TNF receptors on tumor cells, sensitizing them to T-cell-derived TNFα. These findings provide critical mechanistic insights into ADC-immunotherapy combinations and suggest rational strategies for optimizing such regimens in clinical development.

## Results

### Enhanced antitumor activity of ADC and TCE combination

To assess combination effect of ADC and TCE we initially used ADC with TROP2 targeting monoclonal antibody, which was recombinantly produced with complementarity-determining region sequences for antibody clone RS7. ADCs were prepared by conjugation of antibodies to topoisomerase 1 (Top-1i) inhibitor payload (AZ14170133) using maleimide-based linker chemistry(15). T-cell engager contained anti-CD3 and anti-EGFR arms with a silent Fc (16) referred further as TCE. As a target, we used PC9 non-small cell lung cancer (NSCLC) cell line, which express both TROP2 and EGFR on the surface **(Fig. 1A**). PC9 cells were cultured with T cells in the presence of ADC, TCE, or their combination. Antitumor effect was assessed in a Incucyte platform (**Fig. 1B**) or by flow cytometry (**Fig. 1C**). Both assays demonstrated potent augmentation of tumor cell killing when the two drugs were combined. To assess if this effect was dose dependent, we tested in matrix system different concentrations of each agent and found the increase in the killing of tumor cells with the increased concentration of each agent (**Fig. 1D**). This effect was specific for tumor targeting by ADC, since isotype control antibody conjugated with the same payload failed to potentiate the effect of TCE (**Fig. 1E**). Synergy of ADC and TCE agents was evaluated by using highest single agent (HSA) and response additivity (RA) criteria (17). HSA combination index showed that at almost all doses used in the combination, the effect was synergistic (**Fig 1F**). Noticeably, this potent combination effect was observed despite substantial loss of T cells (**Fig. 1G**). To evaluate if ADC-TCE combination had antitumor effect in vivo, we used s.c. tumor model of PC9 in NSG mice humanized with in vitro expanded T cells and then treated with a single dose of ADC and weekly doses of TCE for a total of 3 injections (**Fig. 1H**). Treatment of mice with ADC alone did not (p=0.22) reduce tumor growth. TCE alone caused very modest antitumor effect. In contrast, combination therapy induced profound antitumor activity (**Fig. 1H and Fig. S1**) persisted for at least 20 days (the time of observation). Treatment of mice with TCE was associated with marked increase in the number of tumor-infiltrating T cells (assessed per gram of tissue on day 22 after the start of the experiment) (**Fig. 1I**). However, despite very potent antitumor activity of ADC/TCE combination in comparison to TCE treatment alone, the number of tumor T cells was dramatically lower in combination group than in TCE treated group. Moreover, CD4^+^ T cell number had a trend to be lower when compared with vehicle alone group (**Fig 1I**).

**Figure 1.**
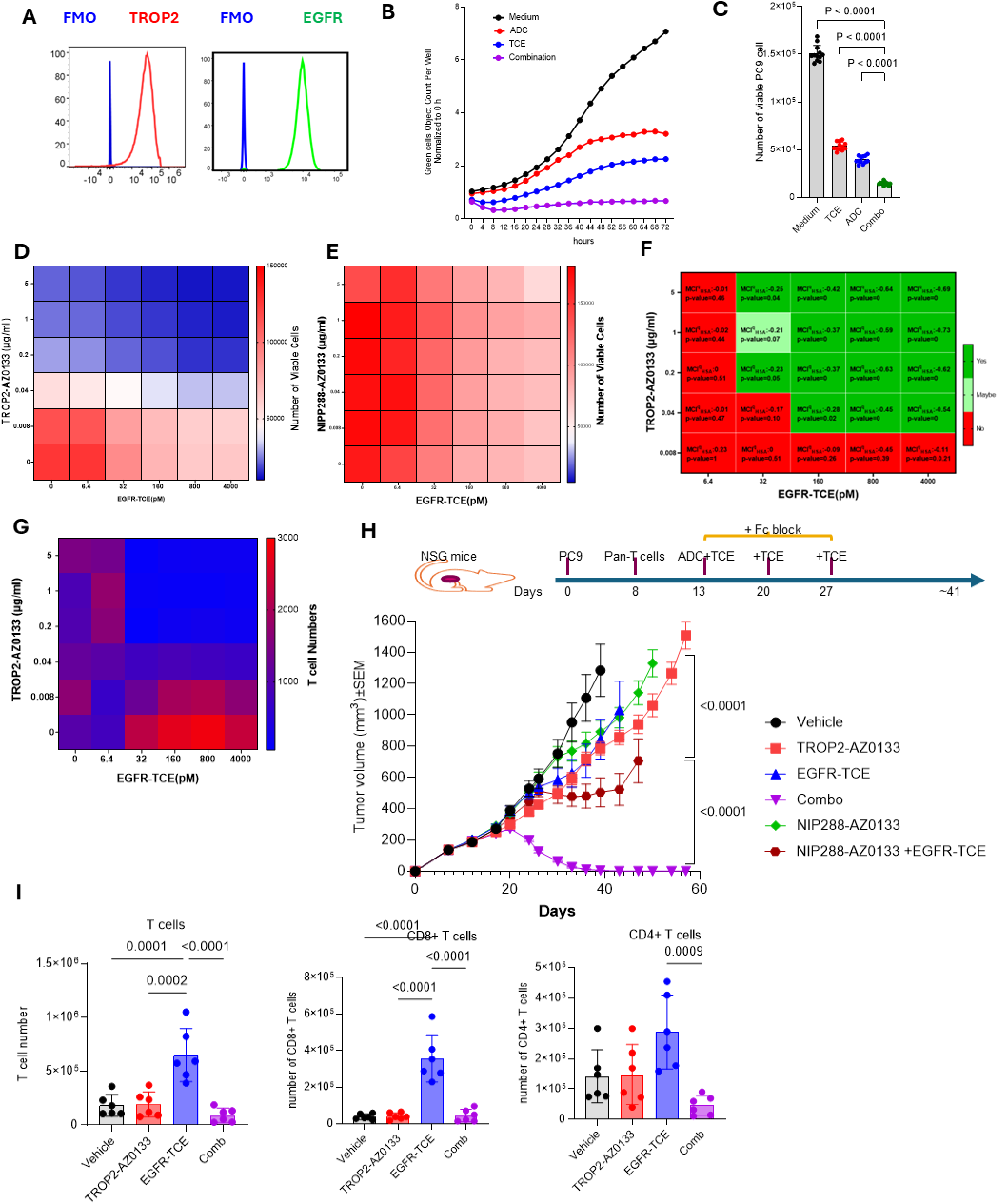
Antitumor effect of ADC and TCE. **A.** Representative flow cytometry plot showing co-expression of TROP2 and EGFR on PC9 tumor cell. **B–C**. Quantification of viable PC9 cells cultured with healthy donor (HD) T cells in the presence of ADC, TCE, or their combination assessed by Incucyte (**B**) or flow cytometry (**C**) (n = 10). P values were calculated in one-way ANOVA with Tukey’s post hoc test. **D.** Heat maps showing viable cancer cell numbers after co-culture with T cells across different TCE (6.4–4000 pM) and ADC (0.008–5 µg/mL) concentrations. **E.** Heat maps showing viable cancer cell numbers after co-culture with T cells across different TCE (6.4–4000 pM) and NIP288–AZ0133 (0.008–5 µg/mL) concentrations. **F.** Heat map of model-based synergy scores quantifying interaction between ADC and TCE (n = 5). HSA synergy index calculated from the cell-killing assay data is shown on the graph. **G**. Viable T-cell counts after co-culture with cancer cells across TCE (6.4–4000 pM) and ADC (0.008–5 µg/mL) concentrations. Viable cell numbers were obtained by absolute counting via flow cytometry (n = 5). **H.** Schematic of the mouse treatment regimen (upper panel) and tumor growth curves (lower panel) for mice treated with ADC, TCE, NIP288–AZ0133 (control), and their combinations (n = 10 per group). P values were calculated in two-way ANOVA test. **I**. Numbers of infiltrating T cells (left), CD8^+^ T cells (middle), and CD4^+^ T cells (right) in tumors from mice treated with ADC, TCE, or their combination. Tumors were collected on the second day after the second dose of TCE. Counts are normalized per gram of tissue. Individual values, mean, and s.d. are shown (n = 6). P values were calculated using unpaired, two-sided Student’s t-tests.

As expected, payload alone caused dramatic decrease in T cell numbers expanded by TCE (**Fig. S2A**). We considered two major mechanisms: a direct effect of ADC on T cells in tumor microenvironment or release of the payload after internalization of ADC by dying tumor cells. Consistent with the data presented in **Fig. 1**, increased concentrations of ADC caused massive drop in TCE induced T cell proliferation and the number of T cells (**Fig S2B**). To test if ADC are directly responsible for T cell killing, experiments were performed in the absence of tumor cells (targets for ADC). T cells were activated by CD3/CD28 since TCE in the absence of tumor targets would not activate T cells. Potent T-cell proliferation and T-cell expansion was observed in response to CD3/CD28 activation. ADC under these conditions failed to inhibit T cell proliferation and decrease the number of T cells (**Fig S2C**) suggesting that only after interaction with tumor cells and release of payload, ADC was toxic for T cells. TCE in addition to T cell activation also provides close proximity of T cells and tumor cells. ADC targeting the same tumor cells would have better accessibility to T cells. To test the importance of this interaction, we used activated T cells with CD3/CD28 in the presence of PC9 tumor cells and ADC. We observed mild decrease in T cell proliferation, but the number of T cells was markedly reduced by ADC (**Fig S2D**). The magnitude of the effect was smaller than in ADC-TCE combination (**Fig.S2B**) suggesting that close proximity of T cells and tumor cells although was not a defining factor, nevertheless contributed to the toxic effect of ADC. These results indicate that ADC mediated loss of T cells depends on ADC interaction with tumor cells and release of payload. Importantly, ADC-TCE combination had potent anti-tumor effect despite substantial reduction in T cells.

### Mechanism of antitumor effect of ADC-TCE combination

Although ADC caused loss of T cells, the remaining cells demonstrated signs of activation. By using concentration matrix of ADC and TCE we found that T cells produced increased amount of IFNγ at the concentrations that caused T cell loss (**Fig. 2A** and **Fig. 1G**).). T-cell surface expression of CD69 activation marker was highest when cells were treated with ADC and TCE (**Fig 2B**). We then evaluated CD8^+^T cells after the combination treatment in vivo by using the same humanized mouse model as in Figure 1H. Expression of 17 surface molecules associated with T-cell function was evaluated in tumor and spleen CD8^+^T cells by flow cytometry. Examples of the analysis are shown in **Fig S3A**. We considered changes as significant if the expression of markers in ADC-TCE combination group was significantly different from both ADC only and TCE only groups. Combination treatment did not demonstrate significant differences in the expression of any of 17 tested markers in spleen CD8^+^ T cells (**Fig. S3B**). In tumor CD8^+^ T cells combination treatment caused substantial up-regulation of CD39, CCR7, CD95 and CD45RO markers – all associated with T cell activation (**Fig. 2D**). These results indicate that ADC-TCE combination can potentiate T cell activation and that effect was confined to tumor site. Next, we isolated tumor infiltrating T cells and tested their ability to kill tumor cells. When T cells were tested directly after isolation their ability to kill tumor cells was the same in mice treated with TCE or ADC-TCE combination (**Fig. 2E**). When those T cells were re-activated *in vitro* with TCE, a modest but significantly increased cytotoxicity was observed by T cells isolated from mice treated with combination therapy (**Fig. 2E**). We also evaluated the effect of the treatment on the cytokine producing CD8^+^ T cells in spleen and tumors. Without in vitro restimulation with TCE, we could not detect spleen or tumor CD8^+^ T cells producing cytokines associate with T-cell activation: IL-2, TNFα, or IFNγ (**Fig. 2F, G**). TCE re-stimulation in vitro dramatically increased the presence of cytokine producing CD8^+^ T cells isolated from spleen (**Fig. 2F**) or tumor (**Fig. 2G**). However, mice treated with combination therapy demonstrated very modest increase in the cytokine producing CD8^+^T cells as compared to mice treated with TCE only (**Fig. 2F, G**). All together these results suggest that activation of T cells can contribute to the effect of combination therapy but is unlikely to be a major factor responsible for the phenomenon of enhanced antitumor activity.

**Figure 2.**
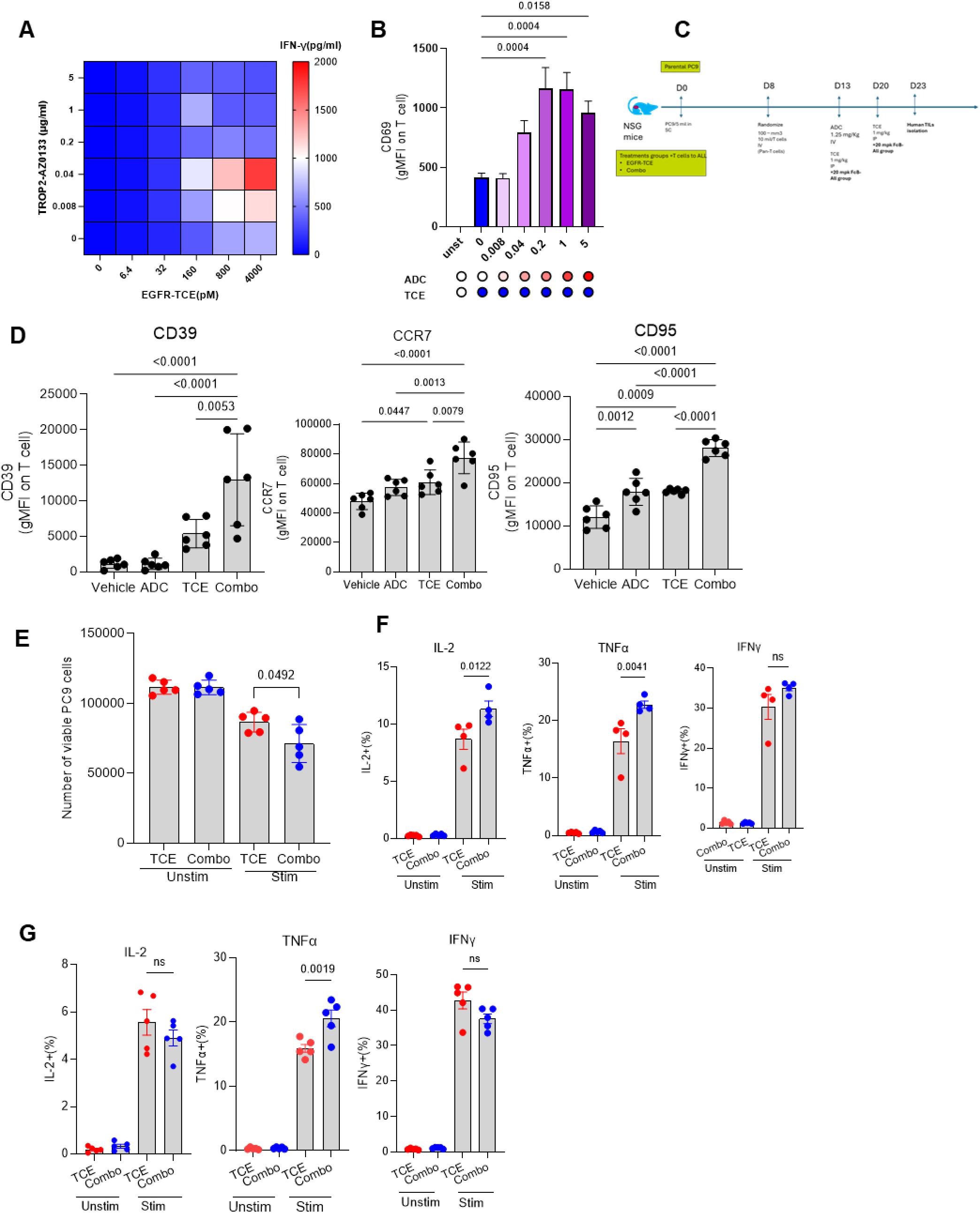
Combination therapy activates T cells. **A.** Heat map quantifying IFN-γ in supernatants from co-cultures of PC9 tumor cells with T cells across different TCE (6.4–4000 pM) and ADC (0.008–5 µg/mL) concentrations. (n = 5). **B.** Expression of the activation marker CD69 on T cells co-cultured with cancer cells in the presence of TCE and ADC (0.008–5 µg/mL) Mean and SD are shown. (n = 5). P values were calculated one-way ANOVA with Tukey’s post hoc test. **C**. Schematic of the in vivo study design showing the treatment schedule and the endpoint for collection of tumors and spleens for T-cell isolation. **D.** Expression of activation markers on tumor-infiltrating T cells from mice treated with TCE, ADC, or their combination. Individual results, mean and SD are shown (n = 5). P values were calculated in one-way ANOVA with Tukey’s post hoc test. **E**. Viable PC9 cancer cell counts in the presence or absence of TCE (160 pM) after co-culture with T cells isolated from tumors of mice treated with TCE alone or the combination with ADC. (n = 5). Individual results, mean and SD are shown. P values were calculated in one-way ANOVA with Tukey’s post hoc test. **F, G.** Percentages of cytokine-positive CD8^+^ T cells (among total CD8^+^ T cells) producing IL-2, TNFα, and IFN-γ. T cells were isolated from tumors (n = 4) **(F)** or spleens (n = 5) (**G)** and re-stimulated ex vivo with a T-cell activation cocktail containing Brefeldin A (n=6). Individual values, mean, and s.d. are shown. P values were calculated in one-way ANOVA with Tukey’s post hoc test.

There are number of factors produced by T cells that can be responsible for their cytotoxic effect. Granzyme B is widely considered as a major mediator of T cell cytotoxicity. We explored the effect of ADC-TCE combination on granzyme B release by T cells. As expected, ADC did not induce Granzyme B release, whereas TCE caused massive release of the enzyme. Combination of ADC and TCE, however, did not change the amount of granzyme B released by T cells (**Fig. 3A**) We previously implicated mannose-6 phosphate receptor M6PR (IGFR2) in antitumor effect of chemotherapy and immunotherapy (18). M6PR can bind granzyme B, that in turn induces the death of the tumor cells. We proposed that a similar mechanism could be involved in ADC-TCE combination. ADC caused considerable up-regulation of M6PR on tumor cells’ surface (**Fig. S4A**). This M6PR upregulation resulted in an increased uptake of recombinant granzyme B by tumor cells (**Fig. S4B**). To assess the functional consequences of M6PR up-regulation, we deleted M6PR in PC9 (**Fig S4C**) and cultured these cells with T in the presence of ADC and TCE. Marked down-regulation of M6PR did not diminish the anti-tumor activity of ADC-TCE combination (**Fig 3B**) indicating that M6PR was not responsible for the observed combination effect of ADC and TCE. Then, we explored other mechanisms that could be involved in T cell mediated killing of tumor cells. We tested blocking antibodies against different molecules involved in T-cell cytotoxicity: TRAIL, IFNAR1, FASL, IFNγ, NKG2D, and HMGB1. For each antibody several dilutions were used based on the highest effective concentrations reported to block their respective ligands (19–25). None of these blocking antibodies were able to abrogate the effect of ADC-TCE combination (**Fig. S5**). In contrast antibody blocking TNFα abrogated the effect of the combination therapy (**Fig 3C**), reverting to the level of ADC or TCE alone. This suggested that in these experimental settings TNFα might be critical for the effect of the combination therapy. TCE but not ADC caused massive release of TNFα. Combination with ADC only slightly increased it (**Fig. 3D**), which was consistent with modest effect of ADC on T cell activation observed in experiments in vitro and in vivo (**Fig. 2**). To validate the effect of T-cell derived soluble factor in this system, we generated supernatants from the culture of PC9 tumor cells with T cells in the presence of TCE. Supernatants then were added to PC9 cells treated with ADC. The supernatants alone did not cause death of tumor cells but potentiated the killing of tumor cells by ADC (**Fig. 3E**). Blockade of TNFα in the medium with antibody abrogated that effect (**Fig 3F**). To confirm the effect of TNFα, we added recombinant TNFα to PC9 tumor cells treated with ADC. TNFα strongly potentiated the antitumor effect of ADC (**Fig 3G**). To determine the effect of TNFα blockade in vivo, mice bearing PC9 tumor were treated with ADC and TCE the same way as described in **Fig. 1H**. Consistent with previous results, ADC-TCE combination caused tumor rejection (**Fig. 4A**). Anti- TNFα antibody did not affect tumor growth when added to untreated mice or mice treated with TCE alone. However, it markedly reduced antitumor effect of ADC-TCE combination (**Fig. 4A**). Thus, TNFα mediated the antitumor effect of ADC-TCE combination.

**Figure 3.**
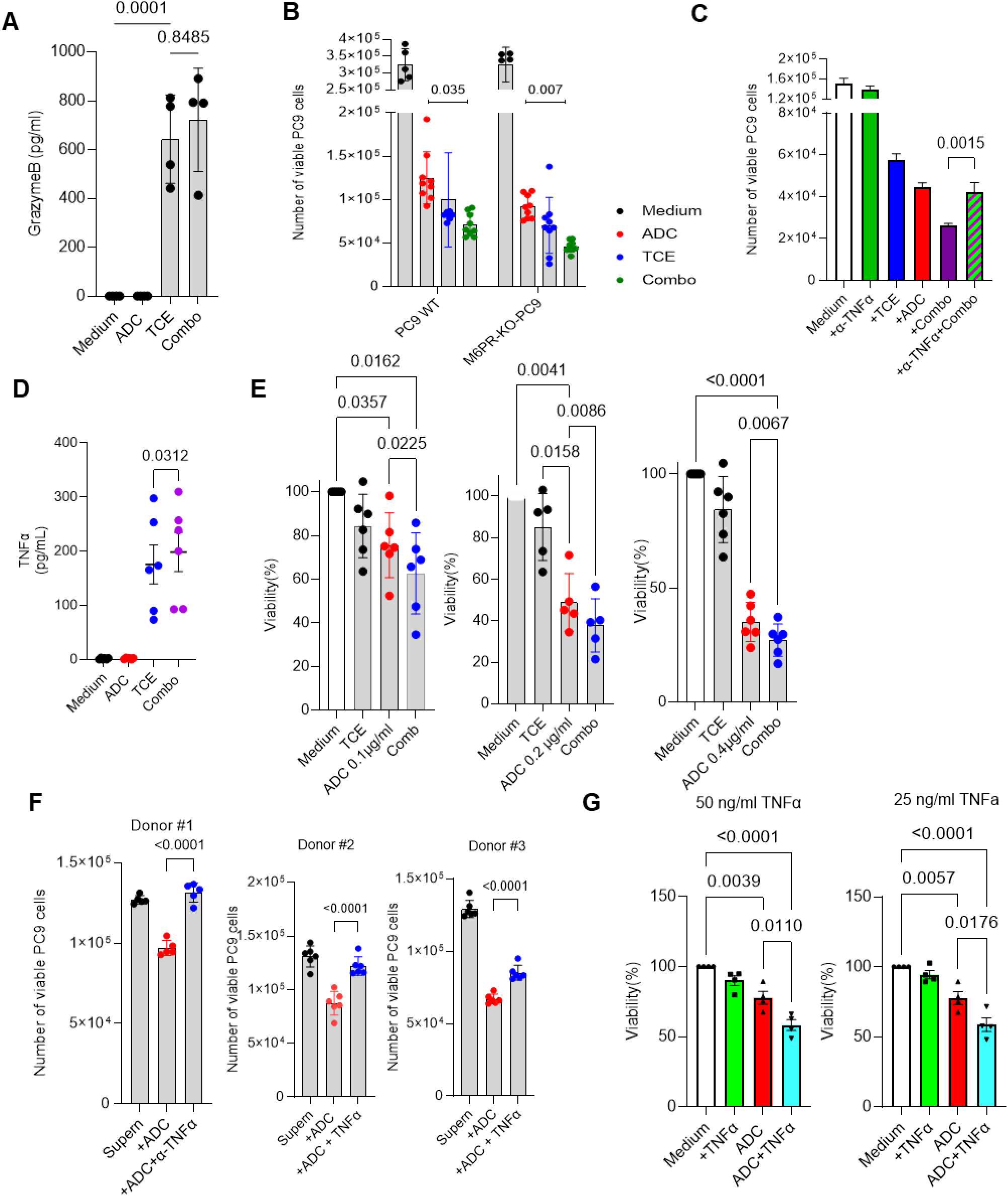
TNFα is the main driver of the effect of ADC-TCE combination. **A.** Granzyme B levels in culture supernatants from PC9 + T-cell co-cultures in the absence or presence of ADC (0.04 µg/ml), TCE (160 pM), or their combination (n=4). **B.** Viable tumor cell counts for PC9 and PC9-M6PR-KO cells after co-culture with T cells in the presence of TCE, ADC, or their combination (n=8). Individual values, mean, and s.d. are shown. P values were calculated using one-way ANOVA with correction for multiple comparisons. **C.** Viable PC9 tumor cell counts after co-culture with T cells with TCE, ADC, or combination in the presence of TNFα antibody. blockade. Mean, and s.d. are shown (n=6). **D**. TNFα concentrations in supernatants from co-cultures of PC9 cells and T cells treated with TCE, ADC, or their combination (n = 6). **D**. Cancer cell viability in the presence of varying concentrations of ADC, supernatants from TCE, or the combination of ADC with TCE supernatants. TCE supernatants were generated by collecting supernatants from PC9 cells co-cultured with T cells in the presence of TCE for 3 days. Assays used a 1:1 mixture of TCE supernatants and fresh medium (n=6). **E**. Cancer cell viability in the presence of ADC and TNFα blockade. N=5. **E**. Cancer cell viability in the presence of ADC, recombinant TNFα (25 or 50 ng/mL), or their combination (n = 3). Unless specified otherwise individual values, mean, and s.d. are shown. P values were calculated in one-way ANOVA with Tukey’s post hoc test.

**Figure 4.**
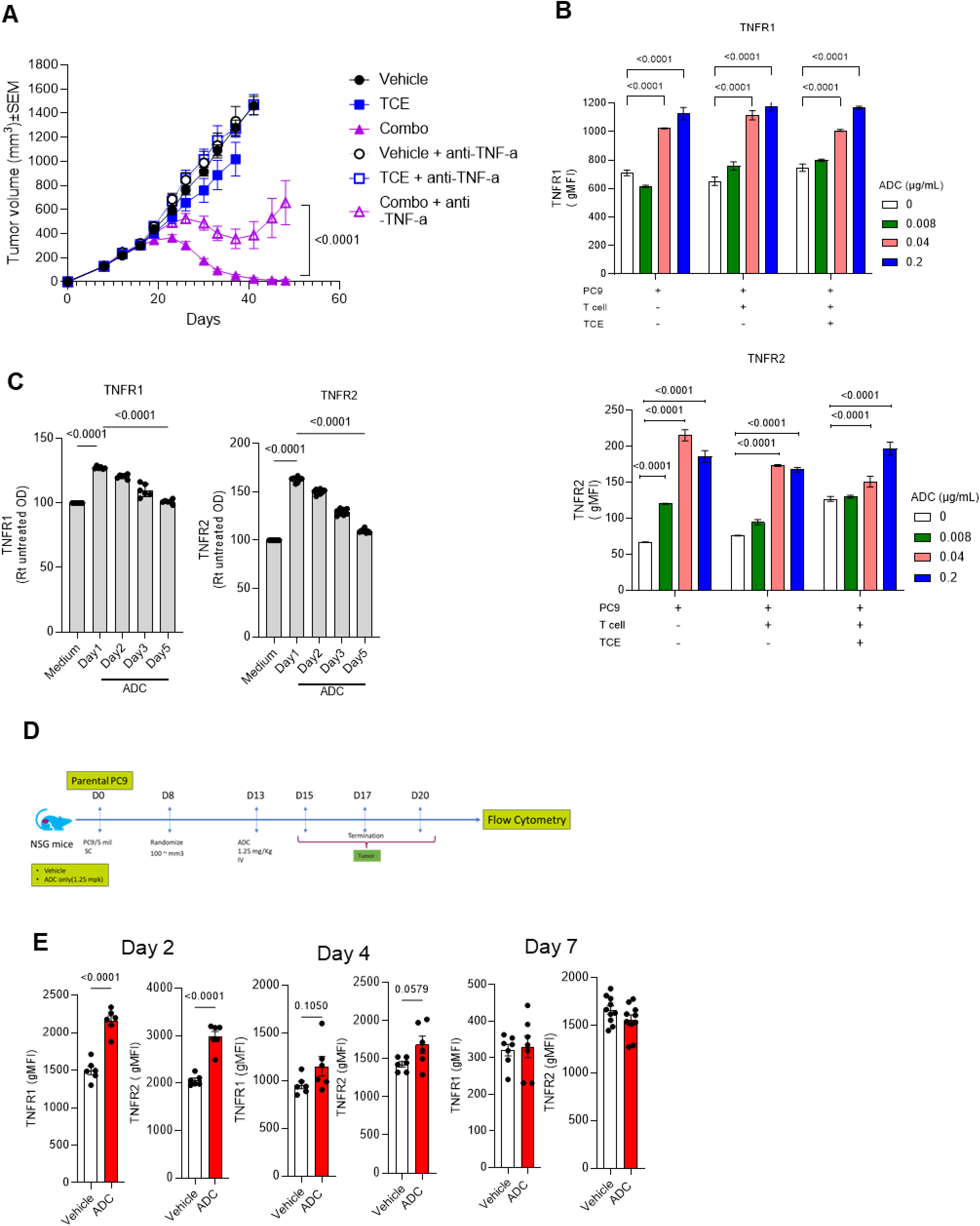
ADC induces upregulation of TNFRs on cancers cells. **A.** Tumor growth curves for mice treated with TCE, the combination of ADC and TCE with or without TNFα blockade with antibody (n = 10 per group). TNFα antibody treatment was initiated 2 days before treatment and continued twice weekly until study ended. **B**. Expression of TNFR1 (left panel) and TNFR2 (right panel) on PC9 cells cultured alone or with T cells in the presence of TCE, ADC, or their combination. N=3. Mean and SD are shown. P values were calculated in one-way ANOVA with Tukey’s post hoc test. **C**. Persistence of TNFR1 (left panel) and TNFR2 (right panel) expression on PC9. Cancer cells were pre-treated with ADC for 3 days to induce upregulation, followed by wash and time-course assessment. N=6. Individual results, mean, and SD are shown. P values were calculated in one-way ANOVA with Tukey’s post hoc test. **D.** Schematic of the in vivo study design, showing the treatment schedule and the endpoint for tumor collection to assess the duration of TNFR1 and TNFR2 upregulation. **E**. Expression of TNFR1 and TNFR2 on tumor cells in vivo following ADC treatment. Panels depict analyses at 2, 4, and 7 days after the first ADC dose. Each point represents individual mouse. N=7. Individual results, mean, and SD are shown. P values were calculated in two-sided Student’s t-tests.

### ADC regulated expression of TNF receptors on tumor cells

Considering the modest increase in TNFα production after ADC-TCE treatment we asked, how ADC could potentiate the antitumor effect of TCE induced TNFα. Treatment of tumor cells with ADC caused substantial dose-dependent up-regulation of TNFR1 and TNFR2 receptors **(Fig. 4B**) suggesting possible mechanism for sensitization of tumor cells to the effect of TCE. The effect was associated only with ADC, since TCE did not affect expression of the receptors (**Fig. 4B**). Although TNFR1 and TNFR2 are both specific receptors for TNFα their biological role is different. TNFR1 is generally associated with inflammation and cell death, while TNFR2 is primarily associated with cell survival, tissue regeneration, and immune regulation (26). TNFRs up-regulation was evident within 24 hr treatment with ADC. After washing, this effect quickly diminished and the expression returned to pre-treatment level by day 4 (**Fig. 4C**). We assessed the changes in TNFRs expression in tumor cells in vivo (**Fig. 4D**). Marked up-regulation of TNFRs was observed 2 days after ADC treatment. The up regulation went down two days later and returned to the pre-treatment level by day 7 (**Fig 4E**). In an attempt to directly evaluate the role TNFRs in antitumor effect of ADC/TCE combination we selectively deleted *TNFR1* in tumor cells (**Fig. S6A**). However, this deletion caused marked increase in the expression of TROP2 and EGFR proteins– targets for ADC and TCE (**Fig. S6B**). As a result, ADC and TCE had substantially better killing on TNFR1 KO cell lines than their parental one (**Fig. S6C**). Thus, the direct assessment of combination effect under those conditions was not possible. To adjust for increased sensitivity of tumor cells to ADC in TNFR1KO tumor cells, we calculated the ratio of tumor cell killing observed after treatment with ADC alone and with ADC-TCE combination. In TNFR1KO cells, combination effect was substantially lower than that in control tumor cells (**Fig. S6D**). To test the effect of ADC on TNFRs expression in patients derived tumors we utilize tumors thick slice technique. Treatment of tissues from breast and lung cancer patients with ADC caused marked up-regulation of TNFR1. TNFR2 up-regulation was significant (p=0.0076) in lung cancer tissues but not breast cancer tissues (**Fig. 5A**). Thus, ADC caused substantial up-regulation of TNFR1 and TNFR2 on tumor cells that could sensitize tumor cells to TCE.

**Figure 5.**
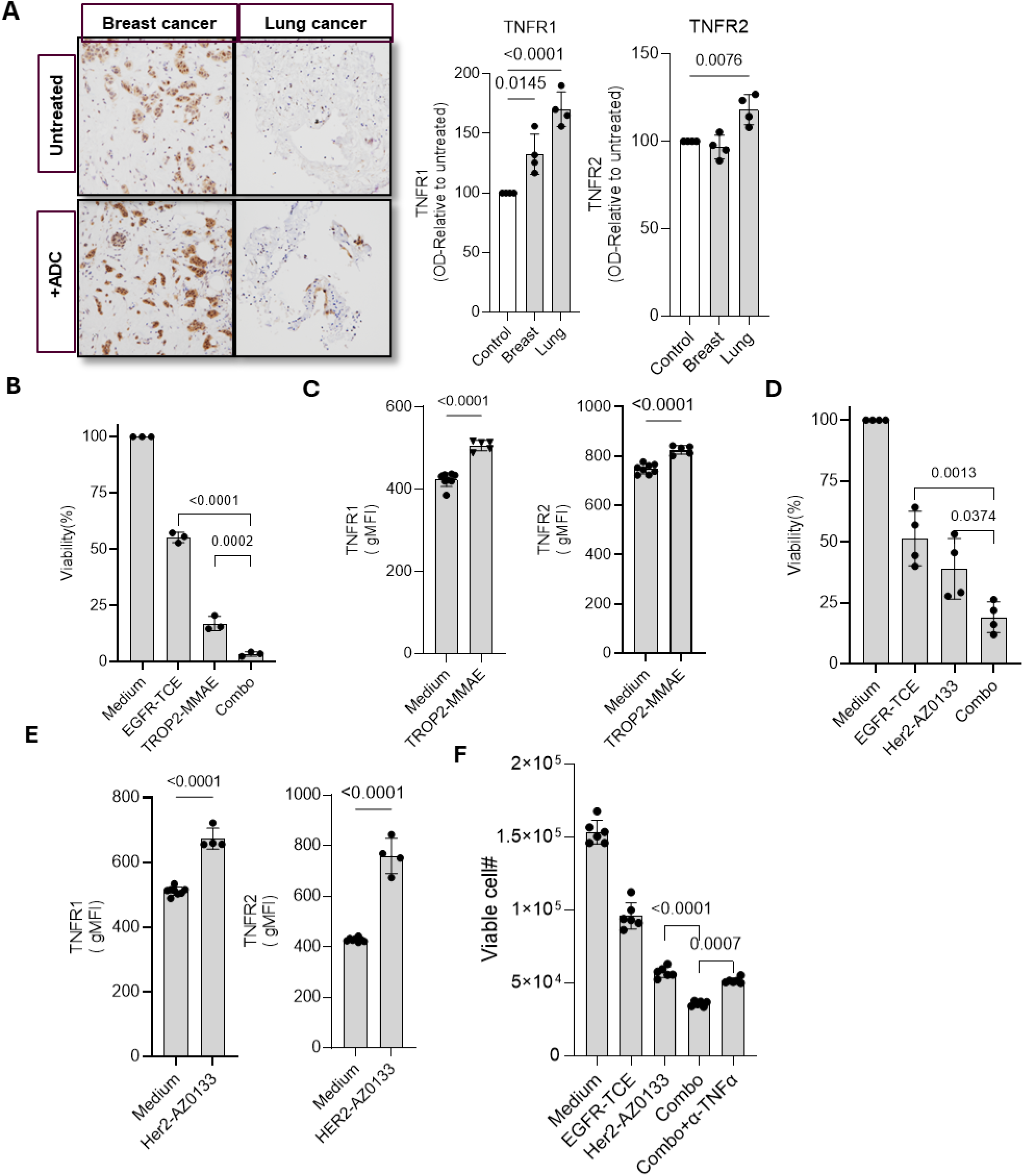
Effect of combination of TCE with various ADC. **A**. Left panels - representative IHC images of TNFR1 staining in patient samples treated with ADC. Right panel -quantification of TNFR1 and TNFR2 IHC staining. Individual values, mean, and s.d. are shown (n = 4). P values were calculated using unpaired, two-sided Student’s t-tests. **B.** Viability of PC9 cells co-cultured with T cells in the presence of TROP2-MMAE, EGFR-TCE, or their combination. Cell numbers were quantified by flow cytometry and normalized to untreated controls (n=3). **C.** Surface expression of TNFR1 (left) and TNFR2 (right) on PC9 cells after treatment with TROP2-MMAE, measured by flow cytometry (n = 5). **D**. Viability of SKBR3 cells co-cultured with T cells in the presence of HER2-AZ0133, EGFR-TCE, or their combination. Cell numbers were quantified by flow cytometry and normalized to untreated controls (n=4). **E**. Surface expression of TNFR1 (left) and TNFR2 (right) on SKBR3 cells after treatment with HER2-AZ0133, measured by flow cytometry (n = 4). **F**. Viable SKBR3 cell numbers in co-culture with T cells in the presence of HER2-AZ0133, EGFR-TCE, or their combination with TNFα blockade. Cell numbers were quantified by flow cytometry (n=5). Individual values, mean, and s.d. are shown. P values were calculated using unpaired, two-sided Student’s t-tests (**C, E)** and one-way ANOVA with correction for multiple comparisons.

### The antitumor effect of ADC-TCE combination was not restricted to specific tumor targets or payload

The experiments described above were all performed with TROP2-Top-1i ADC and TCE combination. We asked if the observed effect was applicable to another payload. We used ADC with monomethyl auristatin E (MMAE), which is a highly potent, synthetic antimitotic cytotoxic ADC payload (27). Combination of TROP2-MMAE ADC with TCE demonstrated potent augmentation of tumor cell killing (**Fig. 5B**). TROP2-MMAE caused up-regulation of TNFRs on tumor cells (**Fig. 5C**) similar to the effect of Top-1i ADC. Next, we used ADC with the Top-1i payload, but targeting a different receptor (HER2) on a different tumor cell line (SKBR3, breast tumor cell line). We observed potent augmentation of antitumor effect in combination with TCE (**Fig. 5D**), which was associated with up-regulation of the expression of TNFRs (**Fig. 5E**). Neutralizing anti-TNFα antibody abrogated combination effect of Her2-AZ0133 with TCE (**Fig. 5F**) recapitulating major findings in other experimental systems.

### ADC induced TNFRs upregulation is mediated by autophagy

Given the role of TNFRs in the effect of ADC-TCE combination we investigated the molecular mechanisms of TNFRs up-regulation by ADC. Treatment with ADC did not cause upregulation of *TNFR1* mRNA but moderate (two-fold) up-regulation of *TNFR2* mRNA (**Fig 6A**). Inhibition of protein synthesis with cycloheximide only marginally affected the surface expression of TNFR1 and TNFR2 (**Fig 6B**) suggesting that changes in transcription and protein synthesis are unlikely to be responsible for ADC induced up-regulation of the receptors on cell surface. Autophagy is associated with the effect of chemotherapy on tumor cells (28,29). We evaluated the possible role of autophagy in up-regulation of TNFRs. ADC treatment of PC9 cells dramatically up regulated lipidated LC3, a hallmark of the autophagy (30) (**Fig 6C**). To directly determine the role of autophagy in the ADC mediated effect, we used two compounds blocking different steps in the autophagy process: bafilomycin, that blocks the vacuolar H^+^ ATPase, necessary for lysosomal acidification and phagosome-lysosome fusion and 3-MA that blocks PI3K activity which is essential for autophagosome formation (31,32). Both compounds abrogated upregulation of TNFR1 and TNFR2 induced by ADC (**Fig. 6D, E**). To validate those observations, we silenced ATG5, the critical component of autophagy with siRNA (**Fig 6F)**. The downregulation of ATG5 substantially reduced TNFR1 and TNFR2 upregulation induced by ADC (**Fig 6G**). Down-regulation of ATG5 abrogated anti-tumor combination effect of ADC and TCE (**Fig. 6H**). Taken together these results suggested that autophagy could be responsible for up-regulation of TNFRs by ADC.

**Figure 6.**
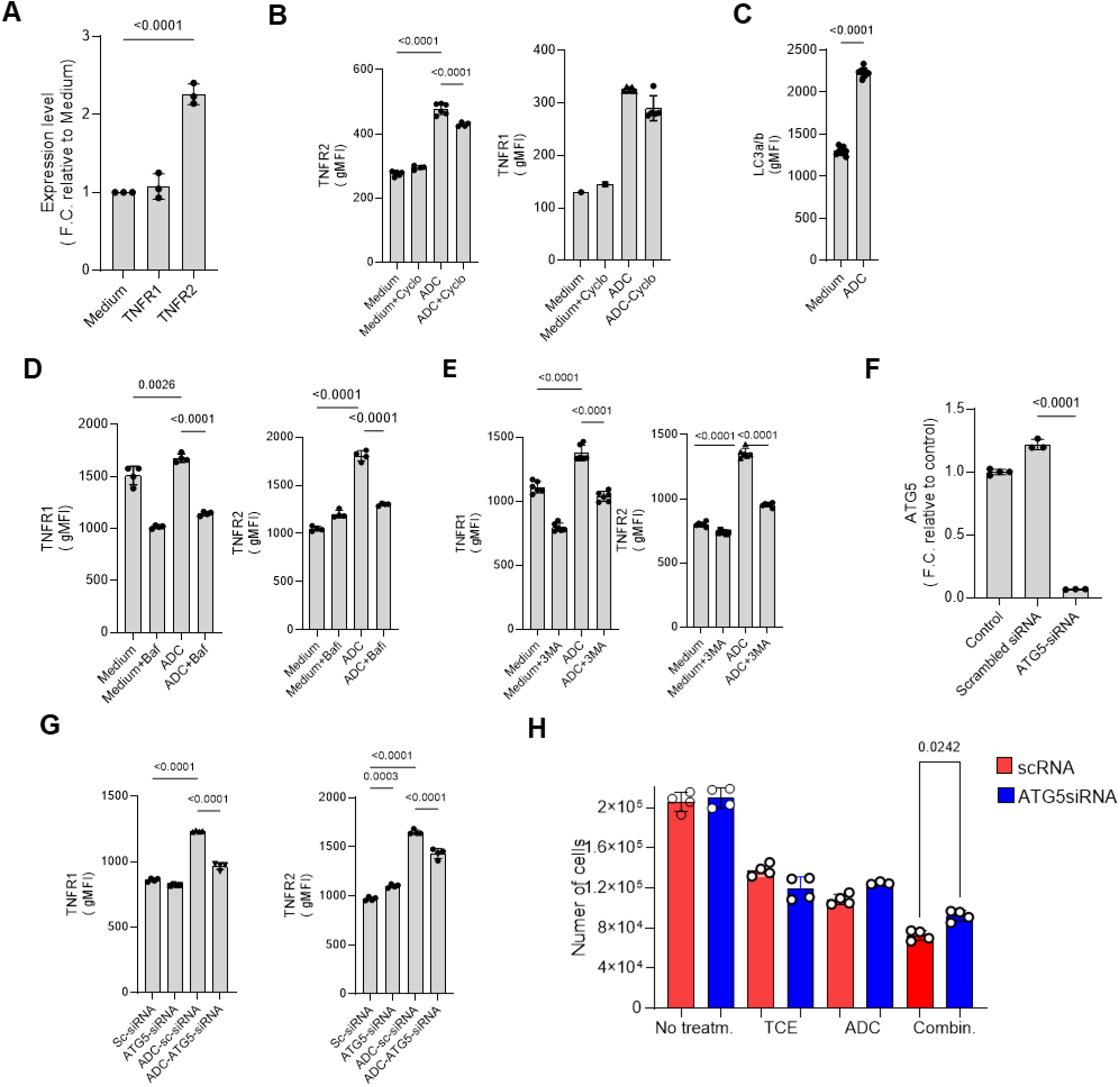
ADC regulates TNF receptors expression through autophagy. **A**. Relative expression of *TNFR1* and *TNFR2* mRNA in PC9 cells treated with ADC (n = 3) measured by qRT-PCR. Individual values, mean, and SD are shown. P values were calculated in one-way ANOVA with Tukey’s post hoc test. **B**. Surface expression of TNFR2 (left) and TNFR1 (right) on cancer cells treated with ADC in the presence of cycloheximide measured by flow cytometry (n=5). Individual values, mean, and SD are shown. P values were calculated in one-way ANOVA with Tukey’s post hoc test. **C**. Median fluorescence intensity (MFI) of intracellular LC3B staining in cancer cells after exposure to ADC. N=9. Individual values, mean, and SD are shown. P values were calculated using unpaired, two-sided Student’s t-tests. **D, E**. Surface expression of TNFR1 and TNFR2 on cancer cells treated with ADC (0.04 µg/ml for 48 hr) in the presence of **(D)** bafilomycin A1 or (**E**) 3-methyladenine (3-MA). Individual values, mean, and SD are shown. N=4. P values were calculated in one-way ANOVA with Tukey’s post hoc test. **F**. Relative expression of *ATG5* in PC9 cells after knockdown with siRNA versus scramble RNA control. Individual values, mean, and SD are shown. N=3. P values were calculated in one-way ANOVA with Tukey’s post hoc test. **G**. Surface expression of TNFR1 and TNFR2 on cancer cells with ATG5 knockdown versus scramble control after ADC treatment (0.04 µg/ml for 48 hr). Individual values, mean, and SD are shown. N=4. P values were calculated in one-way ANOVA with Tukey’s post hoc test. **H**. The numbers of PC9 tumor cells transfected with ATG5 siRNA or scramble siRNA in the presence of ADC, TCE, or their combination. Individual values, mean, and s.d. are shown (n = 4). P values were calculated in one-way ANOVA with Tukey’s post hoc test.

### Mechanism of combination anti-tumor effect of ADC and antigen-specific T cells

Next, we asked whether observed mechanism was specific only for TCE or applicable to a combination of ADC with antigen-specific T cells expanded by check-point inhibitors (CPI). First, we tested the effect of ADC in combination with PD-L1 antibody. We used mouse CT26 tumor expressing human HER2. HER2-ADC with Top-1i payload was administered once a week and PD-L1 antibody twice weekly with two rounds of treatment. Combination of HER2-ADC with PD-L1 antibody markedly enhanced antitumor effect of single agent therapy (**Fig. 7A**). Similar effect was observed in this model when PD-L1 antibody was combined with HER2-MMAE ADC (**Fig. 7B**). We also used a different mouse model - EMT6 tumor expressing human HER2 (**Fig. S7A**). Mice were treated with Anti-HER2-Top-1i ADC in combination with PD-L1 antibody. Combination treatment resulted in marked increase in antitumor effect (**Fig. S7B**).

**Figure 7.**
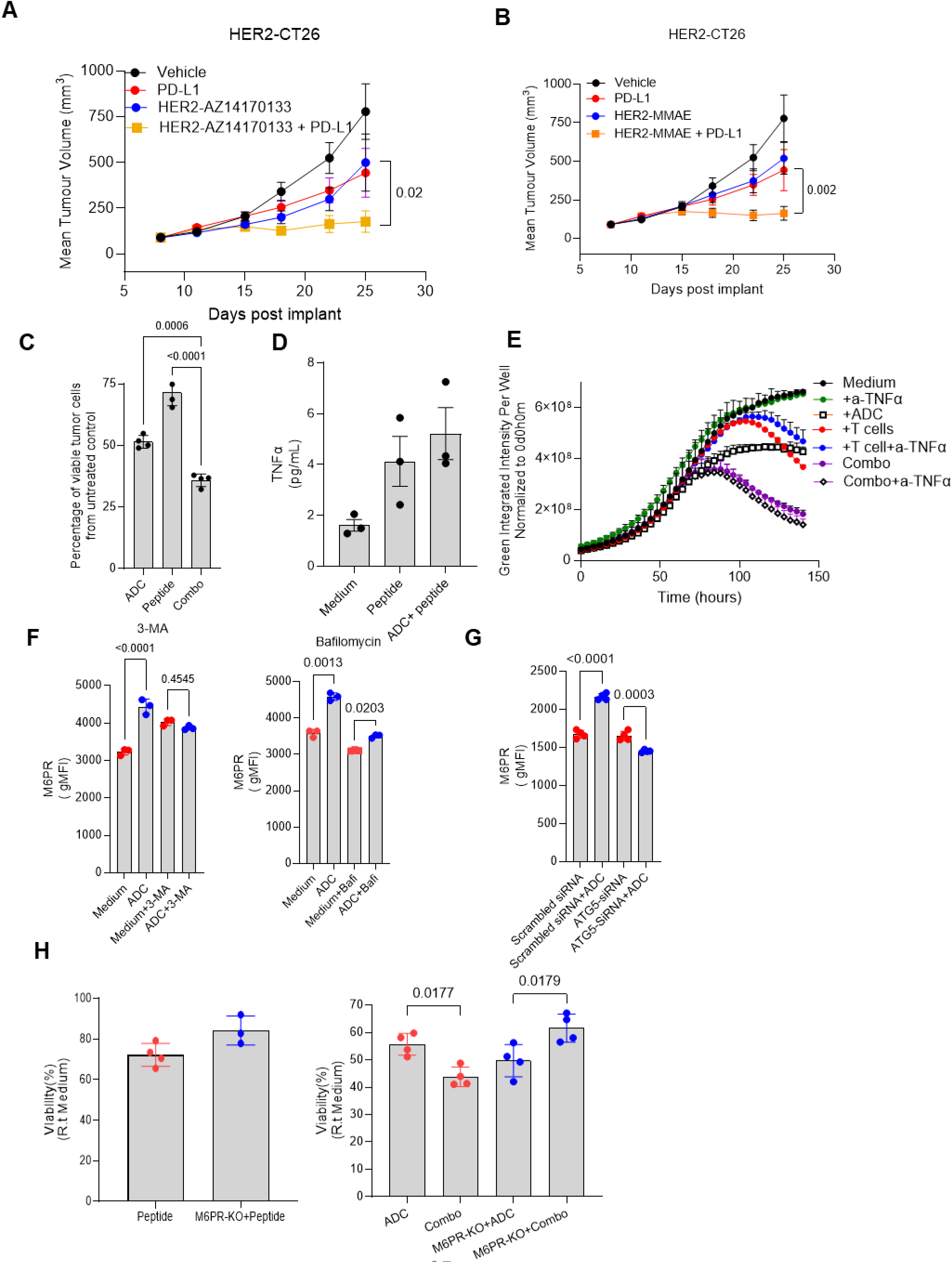
Mechanism of ADC combination to antigen-specific T cells. **A, B.** Growth of CT-26 tumor expressing HER2 in mice treated with PD-L1 and HER2-ADC with Top-1i payload (**A**), or HER2-ADC with MMAE payload (**B**). ADCs were administered once a week and PD-L1 antibody twice weekly with two rounds of treatment. Each group includes 6 mice. Mean and SD is shown. P values were calculated in two-way ANOVA. **C.** Proportion of live PC9 tumor cells loaded with MART1 derived peptide cultured with MART1 specific T cells in the presence of TROP2-Top-1i ADC. Individual values, mean, and SD are shown. N=4. P values were calculated in one-way ANOVA with Tukey’s post hoc test. **D.** The mount of TNFα produced by peptide-specific T cells treated with ADC. Individual values, mean, and SD are shown. N=4. **E.** Live PC9 tumor cells loaded with MART1 peptide treated with antigen-specific T cell in the presence of ADC and anti-TNFα antibody. **F.** Effect of 3-MA and bafilomycin on ADC induced up-regulation of M6PR in PC9 tumor cells. Individual values, mean, and SD are shown. N=3. P values were calculated in one-way ANOVA with Tukey’s post hoc test. **G.** Effect of ATG5 deletion in PC9 tumor cells on ADC induced up-regulation of M6PR measured by flow cytometry. Individual values, mean, and SD are shown. N=4. P values were calculated in one-way ANOVA with Tukey’s post hoc test. **H.** Proportion of viable MART1 peptide-loaded M6PR deficient and wild-type tumor cells PC9 tumor cells after treatment with antigen-specific T cells alone (left panel), ADC alone and combination of T cells and ADC (right panel). Individual values, mean, and SD are shown. N=4. P values were calculated in one-way ANOVA with Tukey’s post hoc test.

CPI expand tumor antigen specific T cells. To better understand the mechanism of the observed effect of CPI combination with ADC, we generated MART1 peptide specific T cells by stimulation of donor’s T cells with peptide loaded DCs. This generated 70% of MART1 dextramer positive CD8^+^ T cells (**Fig. S7C**). MART1 specific T cells were cultured with PC9 cells loaded with MART1 peptide in the presence of TROP2-Top-1iADC. Similar to the effect observed with TCE, combination of antigen-specific T cells with ADC potentiated the effect of ADC treatment alone (**Fig. 7C**). In contrast to TCE, peptide specific T cells produced negligible (single pg/ml concentrations) (**Fig. 7D**). Consistent with these results neutralizing TNFα antibody failed to reduce antitumor effect of ADC and antigen-specific T cells combination (**Fig. 7E**).

ADC, in addition to TNFRs, caused up-regulation of M6PR (**Fig. S4)**. We assessed the changes in M6PR expression on tumor cells in mice in response to ADC treatment in vivo. Significant (p=0.0009) up-regulation of M6PR expression was observed in tumor cells 2 days after ADC administration. It remained significantly (p=0.0002) up-regulated 4 days after ADC administration (**Fig. S7D**).

We investigated whether autophagy was responsible for this phenomenon. Autophagy inhibitor 3-MA abrogated ADC induced up-regulation of M6PR and bafilomycin markedly reduced the effect (**Fig. 7F**). Deletion of ATG5 in tumor cells completely abrogated ADC induced up-regulation of M6PR (**Fig. 7G**). Thus, autophagy was responsible for ADC mediated up-regulation of M6PR. We tested if deletion of M6PR could reverse combination effect of ADC and antigen-specific T cells. Deletion of M6PR in tumor cells did not affect killing of cell by ADC or T cells (**Fig. 7H**). However, enhanced killing of tumor cells by a combination of ADC and T cells was abrogated in M6PR KO cells (**Fig. 7H**).

## Discussion

In this study, we report on a mechanism by which ADCs may potentiate T-cell-mediated antitumor immunity through autophagy-dependent upregulation of TNFR and M6PR on cancer cells. Our findings demonstrate that combining ADCs with TCE results in synergistic tumor cell killing despite substantial depletion of tumor-infiltrating lymphocytes. We, for the first time, report antitumor effect of ADC-TCE combination in solid tumors and identified TNFα-TNFR signaling as the critical mediator of this enhanced cytotoxicity in the ADC-TCE setting, while revealing a distinct M6PR-dependent mechanism operative when ADCs are combined with CPI expanded antigen-specific T cells (**Fig. S8**).

The demonstration of potent synergy between ADC and TCE, as quantified by HSA analysis (33,34), was paradoxical given the concurrent dramatic reduction in T-cell numbers. This apparent contradiction challenged conventional assumptions that greater T-cell infiltration is required for superior immunotherapeutic efficacy and suggested that the remaining T cells possessed enhanced cytotoxic capacity or that ADC treatment fundamentally altered tumor cell susceptibility to T-cell-mediated killing. Granzyme B, widely regarded as the primary effector molecule of cytotoxic T lymphocytes (35), showed no differential release in the combination setting compared to TCE alone. Similarly, blocking antibodies against multiple immune effector molecules failed to abrogate the combination effect. In contrast, TNFα blockade completely reversed the enhanced cytotoxicity to levels observed with single agents, establishing TNFα as possible essential mediator of ADC-TCE synergy. TCE induced release of large amount of TNFα by T cells, which was only minimally affected by addition of ADC. Instead, ADCs substantially upregulated both TNFR1 and TNFR2 expression on tumor cell surfaces in a dose-dependent manner. TNFR1 contains a death domain and primarily mediates pro-apoptotic and pro-inflammatory signaling, whereas TNFR2, which lacks a death domain can contribute to cell death under certain contexts (36). This receptor upregulation effectively sensitized tumor cells to TNFα produced by TCE-activated T cells. The functional validation through experiments using TCE-conditioned supernatants and recombinant TNFα as well as in vivo validation with anti-TNFα blocking antibodies confirmed that ADC-treated tumor cells became sensitive to TNFα-mediated killing. Partial rescue of antitumor effect in vivo with TNFα antibody suggests that additional mechanisms may contribute to combination activity, or that incomplete TNFα neutralization was achieved in the tumor microenvironment. This results were consistent with recent observation that TNFα potentiated the effect of TCE mediated tumor cell killing (37)

Autophagy, a highly conserved cellular process involving the degradation and recycling of cytoplasmic components through lysosomal pathways, is a critical mediator of cancer cell responses to chemotherapy and targeted agents (28,29). Our finding that ADC treatment induced robust LC3 lipidation, a hallmark of autophagosome formation (38), confirmed that ADC induces autophagy in tumor cells. Autophagy was responsible for ADC mediated up-regulation of TNFR and M6PR on tumor cells. Autophagy can modulate TNFR and M6PR trafficking to the cell surface rather than inducing new protein synthesis. The kinetics of TNFR upregulation—evident within 24 hours of ADC treatment and reversible upon drug removal—indicate a dynamic regulatory process rather than stable epigenetic changes.

A particularly noteworthy observation is that the mechanism of ADC-immunotherapy synergy differs depending on the mode of T-cell activation. While ADC-TCE combinations depended on TNFα-TNFR signaling, combinations with antigen-specific T cells operated through a TNFα-independent, M6PR-dependent pathway. This mechanistic dichotomy likely reflects fundamental differences in T-cell activation states and effector molecule production between TCE and antigen-specifically activated T cells. TCE binding to CD3 triggers robust T-cell activation and cytokine production, including copious TNFα secretion. In contrast, peptide-specific T cells, while effectively killing target cells presenting cognate antigen, produced minimal TNFα. Instead, the ADC-mediated antitumor effect in this setting required M6PR (IGF2R), a multifunctional receptor involved in lysosomal enzyme trafficking and growth factor signaling. Previous work has implicated M6PR in granzyme B uptake that mediate combination effect of T-cell therapy and targeted therapy or chemotherapy of cancer (18,39,40). Our finding that ADCs upregulate M6PR expression through an autophagy-dependent mechanism, identical to the pathway regulating TNFRs, reveals a common upstream driver (autophagy) that modulates multiple surface receptors relevant to immune-mediated cytotoxicity (**Fig. S8**). The evaluation across different variables (alternative payloads, different target antigens, distinct tumor types, and various immunotherapy modalities) demonstrated remarkable consistency in the core mechanism. Autophagic response represents a common cellular stress pathway activated by diverse cytotoxic insults rather than a payload-specific effect.

Our findings suggest that concerns about payload-mediated immunosuppression, while valid, may be outweighed by enhanced tumor cell susceptibility to immune attack. However, the optimal dosing and sequencing remain to be determined. The identification of TNFR and M6PR upregulation as key mediators suggests these receptors as potential pharmacodynamic biomarkers. Serial tumor biopsies assessing TNFR expression before and after ADC treatment could provide proof-of-mechanism in early-phase clinical trials and potentially identify patients most likely to benefit from combination therapy. Baseline TNFR expression levels might also serve as predictive biomarkers, with tumors expressing lower levels potentially showing greater dynamic range for ADC-induced sensitization.

While our data demonstrate that autophagy is required for the combination benefit, clinical autophagy inhibitors (such as hydroxychloroquine) should be avoided in combination with ADC-immunotherapy regimens, as they would be predicted to block the beneficial sensitization effect. The mechanistic differences between TCE-based and antigen-specific T-cell-based combinations suggest that biomarker strategies and optimization approaches may need to be tailored to the specific immunotherapy modality.

### Limitations of the study

Several limitations of our study warrant acknowledgment. First, our in vivo experiments utilized immunodeficient mice humanized with in vitro-expanded human T cells, which incompletely recapitulates the complex immune microenvironment of spontaneous human tumors. Currently there are no syngeneic mouse models that adequately recapitulate the effect of ADC and TCE. Future studies with the development of fully immunocompetent mouse models would provide more comprehensive assessment of immune interactions. Second, our mechanistic studies focused on early events occurring within days of combination treatment, whereas durable antitumor immunity often develops over weeks to months. Long-term studies assessing immunological memory, epitope spreading, and durability of responses would provide important additional insights. Third, the question of optimal combination dosing and scheduling requires systematic study.

## Methods

### Patients sample preparation and staining

The study is being performed in accordance with consensus ethical principles derived from international guidelines including the Declaration of Helsinki and Council for International Organizations of Medical Sciences (CIOMS) International Ethical Guidelines, applicable International Council for Harmonization (ICH) Good Clinical Practice (GCP) Guidelines, all applicable laws and regulations, and the AstraZeneca policy on Bioethics and Human Biological Samples. Studies were approved by Institutional Review Board of AstraZeneca. Peripheral blood was collected from healthy volunteers at AstraZeneca after obtaining informed consent.

De-identified patient samples were procured through the National Disease Research Interchange (NDRI). Fresh lung and breast tumor tissues were obtained from patients undergoing surgical procedures. After resection, left over tumor tissue designated for research purposes were kept at 4°C and transported in fresh DMEM + antibiotics. Specimens arrived at the AstraZeneca laboratory within 24 hours of surgery. Section were embedded in 6% agarose and automated slicing was performed using a Leica VT 1200S Vibratome with slice thickness set at 300 µm. Tumor slices were transferred to organotypic inserts and incubated on a plate shaker in a humidified incubator at 37°C, 5 % CO2 in the appropriate volume of complete culture media and plate format (lung tumor tissue: 30 mm insert/6-well plate/1.1 mL media per well, breast tumor tissue: 12 mm insert/24-well plate/0.3 mL media). After 2 hours, media was replaced with appropriate fresh complete culture media alone or + 20 nM ADC. Tumor slices were fixed in 10% neutral buffered formalin for at least 24 hours at room temperature. Subsequently, tumor slices were embedded in paraffin and 4 μm sections were generated for microscopy analysis.

#### TNFR-1 staining

Santa Cruz Biotechnology Inc. TNF-R1 mouse monoclonal IgG2b antibody (H-5) (Cat. sc-8436) was used for immunohistochemistry (IHC). The IHC antibody was diluted to a working concentration of 2µg/ml in Dako/Agilent diluent (Cat. S3022). The fully automated IHC protocol was performed on the Leica Bond Rx (Leica Biosystems Buffalo Grove, IL, USA). Antigen retrieval was performed with BOND Epitope Retrieval Solution 2 (pH 9, Cat. AR9640) for 20 minutes at 100°C. Antibody detection was performed with a Leica Bond Polymer Refine Detection kit: post primary reagent (Cat. DS9800).

#### TNFR-2, M6PR staining

Signaling Technology TNF-R2 (E8D7P) Rabbit Monoclonal Antibody #72337 was diluted to 1µg/ml. Cell Signaling Technology IGF-II Receptor/CI-M6PR (D8Z3J) Rabbit Monoclonal Antibody #15128 was diluted to 0.5µg/ml. Antibody detection was performed with a Leica Bond Polymer Refine Detection kit, using only the polymer reagent for 8 minutes and visualized with 3,3’ diaminobenzidine (DAB) (Cat. DS9800).

### Mouse Models

Mouse experiments were approved by the Institutional Animal Care and Use Committee of AstraZeneca (Gaithersburg, MD) and conducted in Association for Assessment and Accreditation of Laboratory Animal Care (AAALAC)–accredited and United States Department of Agriculture (USDA)–licensed facility. All mice were housed in autoclaved cages with access to food and water *ad libitum* in a sterile environment maintained with a 12hr dark/12hr light cycle at 72±2°F with 50±20% room humidity. Female 8- to 10-week-old NSG (NOD.Cg-Prkdcscid Il2rgtm1Wjl/SzJ) mice were obtained from The Jackson Laboratory.

### Tumor Cell Lines

PC9 non-small cell lung cancer, SK-BR-3 hypertriploid human breast cancer, and murine EMT6 cell lines were obtained from ATCC. PC9 cells were cultured in RPMI 1640 (Gibco) with 10% FBS at 37°C, 5% CO_2_. SK-BR-3 cells were cultured in McCoy’s 5A (Gibco) supplemented with 10% FBS. EMT6 cells were cultured in Waymouth’s MB 752/1 medium with 2 mM L-glutamine and 15% fetal bovine serum. Unless otherwise stated, NSG (NOD.Cg-Prkdcscid Il2rgtm1Wjl/SzJ) mice were subcutaneously injected with 5 × 10^6^ PC9 tumor cells in sterile 1× DPBS into the right flank using a 27.5 G needle on Day 0 of the study. Tumors were measured with digital calipers twice weekly until the end of the study or until they reached the ethical limit of 2,000 mm^3^.

### In Vivo Treatment

Unless otherwise stated, NSG (NOD.Cg-Prkdcscid Il2rgtm1Wjl/SzJ) mice were subcutaneously injected with 5×10^6^ PC9 tumor cells in sterile 1x DPBS into the right flank using a 27 ½g needle on Day 0 of study. Tumors were measured using digital calipers measured twice a week until the end of study or when tumors reached the ethical limit of 2000mm^3^. Mice were randomized and treated when tumors reached 100mm^3^. Mice were given 10^6^ T cells intravenously via the lateral tail vein. T cells were isolated and expanded for 8 days using the Human Pan T Cell Isolation Kit (Miltenyi Biotec, catalog no. 130-096-535) following the manufacturer’s instructions. Unless otherwise indicated, ADC was given once. ADC was diluted in sterile 1x DPBS and administered 200µL intravenously at 1.25mg/kg on Day 13 post tumor implantation. All mice were given intraperitoneally 200µL of anti-mouse CD16/CD32 blocking antibody (BioXCell, Clone 2.4G2) at 20mg/kg prior to TCE treatment to reduce Fcγ receptor–mediated interactions. TCE was diluted in sterile 1x DPBS and administered 200µL intraperitoneally at 0.125mg/kg on Day 13, Day 20, and day 27 post tumor implantation. For studies looking at T cell function and profile, the dosing schedule indicated above was followed, mice were euthanized via CO_2_ asphyxiation, then whole tumors and spleens were harvested on Day 22 or Day 23. For studies of TNFα impact on combination treatments, mice were given 200µL intraperitoneally of anti-TNFα (BioXCell, Infliximab) at 10mg/kg on Day 11 and Day 13 post tumor implantation. Tumor volume was calculated as V = 0.5xLengthxWidth^2^.

### EMT6-hHER2 cell line generation

A single-cell EMT6-hHER2 line was generated by lentiviral transduction of human HER2 under the EF1α promoter. Lentiviral particles were packaged in HEK293T cells using pKLV2-EF1α-HER2, psPAX2, and pMD2.G plasmids. Cells were stained with anti-human CD340 (ErbB2/HER2) Alexa Fluor 488–conjugated antibody (BioLegend, Cat. 324410), and single-cell clones were isolated.

### Human T cell isolation

Human T cells were isolated from peripheral blood mononuclear cells (PBMCs) by negative selection using the EasySep™ Human T Cell Enrichment Kit (STEMCELL Technologies, cat. #19051) according to the manufacturer’s instructions. Purified T cells were washed and resuspended in the appropriate assay medium prior to co-culture.

### Assessment of combination effect of ADC and TCE

To generate heat maps of drug synergy and T cell–mediated cytotoxicity, cancer cells were co-cultured with isolated T cells in the presence or absence of varying concentrations of ADC and TCE for three days. Optimal concentrations determined by these matrices were used for subsequent studies. Unless otherwise specified, 0.04 μg/mL of ADC and 160 pM TCE were used for combination studies. The same workflow was applied to evaluate TROP2-MMAE and HER2-AZ0133 ADCs. To probe mechanisms underlying observed synergy, blocking antibodies—anti-TNFα (R&D Systems, cat. #MAB210), α-TRAIL (R&D Systems, cat. #MAB375), α-IFNAR1 (anifrolumab), α-FasL (R&D Systems, cat. #AB126), α-IFNγ (InvivoGen, cat. #hifng-mab7-02), α-NKG2D (R&D Systems, cat. #MAB139), and α-HMGB1 (BioLegend, cat. #651414)—were added immediately after combining T cells and cancer cells, followed by addition of the ADC and TCE.

After three days of co-culture, plates were centrifuged and supernatants were collected for cytokine analysis. Adherent cells were detached with TrypLE (Gibco, cat# #12605010) after 20 minutes of incubation. Plates were washed with PBS containing 1% FBS and 2 mM EDTA, then with PBS alone. Cells were stained with a live/dead viability dye and surface markers to distinguish cancer cells from T cells and to assess T cell activation. Absolute viable cell counts were obtained by flow cytometry using CountBright Absolute Counting Beads (ThermoFisher, cat# C36950), according to the manufacturer’s formula: exact number of cells = (recorded cell events / recorded bead events) × number of beads added. Samples were acquired on a BD flow cytometer (BD FACSymphony A3)

### Cytokine Measurements

After 3 days of co-culture, supernatants were collected to measure levels of IFN-γ and TNF-α by using the IFN-γ Human ELISA Kit (Thermo Fisher, Cat# KHC4021) and the TNF-α Human ELISA Kit (Thermo Fisher, Cat# KHC3011). Absorbance was read at 450 nm on Spectramax M5 plate reader (Molecular Devices). Cytokine concentrations were determined from a standard curve generated using appropriate curve-fitting software.

### Data acquisition and analysis

Samples were acquired on a Cytek Aurora I-5L spectral flow cytometer equipped with UV (355 nm), violet (405 nm), blue (488 nm), yellow-green (561 nm), and red (640 nm). Data were analyzed in FlowJo v10.10.0 (BD). For dimensionality reduction, tSNE was run on arcsinh-transformed marker intensities using default parameters (perplexity 30, theta 0.5, iterations =1000). Clustering was performed with FlowSOM in FlowJo on the same transformed features, using a 10×10 self-organizing map with hierarchical metaclustering.

This study included five donors. Cells from each donor were treated with ADC and TCE at dose pairs (*d_i_^a^, d_j_^t^*) where *i* = 0, …,5 and *j* = 0, …, 5; here, *d_i_^a^* and *d_j_^t^* denote the ADC and TCE doses, respectively. Control conditions were defined as follows: (*d*_0_*^a^, d*_0_*^t^*) = (0,0) for the untreated control, (*d*_0_*^a^, d_j_^t^*) = (0,*d_j_^t^*) with *j* > 0 for TCE monotherapy, and (*d_i_^a^, d*_0_*^t^*) = (*d_i_^a^*,0) with *i* > 0 for ADC monotherapy. Accordingly, the study employed a repeated-measures design, with multiple dose combinations administered to each donor. For each donor, up to two technical replicates were performed to measure cytotoxicity (killing) and IFNγ responses. When two replicates were available, their mean was used as the recorded value for analysis.

The objective is to assess whether the combination therapy exhibits synergy relative to the two monotherapies with respect to the killing endpoint. Specifically, for each dose pair (*d_i_^a^, d_j_^t^*) with *i* = 1 ⋯ 5, *j* = 1 ⋯ 5, we test:

- *H*_0_: the combination therapy demonstrates no synergy
- *H_a_*: the combination therapy demonstrates synergy

Synergy occurs when a combination produces an effect greater than the simple additive expectation derived from each drug’s individual contribution. Translating this concept into a rigorous quantitative framework is non-trivial; over the past few decades, numerous approaches have been proposed, broadly categorized into effect-based and dose-based methods (34). Dose-based approaches are often considered more precise for quantifying synergism, but they impose stringent data requirements—namely, well-characterized full dose–response curves for each agent alone and in combination—and they rely on complex mathematical inverse operations applied to those curves. These constraints make dose-based methods difficult to apply to our dataset. Accordingly, we adopt effect-based approaches, which are more tractable to compute and more straightforward to interpret given the structure of our current data. Two definitions of synergy are considered: Response Additivity and Highest Single Agent (HSA). The analysis proceeds hierarchically. First, additivity is evaluated; if the additivity criterion is not satisfied, the assessment then proceeds to the HSA framework (17).

The HSA criterion declares synergy when the combination effect exceeds the best single agent, i.e., *E_ij_* > max{*E_i_*_0*k*_, *E*_0*jk*_}, while the RA criterion declares synergy when the combination exceeds the additive expectation, i.e., *E_ijk_*> *E_i_*_0*k*_ + *E*_0*jk*_.

The corresponding HSA and RA combination indexes(17) can be defined as

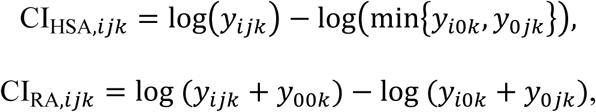

where negative values indicate combination performance superior to the respective monotherapies (consistent with synergy) under the convention that lower *y* reflects greater killing.

For each donor and each synergy definition, the 25 combination indices (CIs) inherit the repeated-measures structure and are mutually correlated. This correlation arises because all CIs for a given donor share common monotherapy and untreated benchmarks and are derived from measurements taken within the same individual.

### Granzyme measurement in supernatant

After 3 days of co-culture, supernatants were collected to measure levels of granzyme B in cell culture. Luminex bead-based customized immunoassay plates (PROCARTAPLEX 10 PLEX, Thermofisher, D5370612) were used for detection according to the manufacturer’s instructions.

### Real-time killing assay

To confirm synergy between ADCs and TCEs using orthogonal readouts, live-cell imaging was performed with the Incucyte system. Cancer cells were labeled with a CellTracker Green CMFDA Dye(Thermo Fisher, Cat#C7025) and co-cultured with T cells in the presence or absence of ADCs. Four images per well were captured at each time point and used to quantify total cell counts. Cell numbers were normalized to the 4-hour time point after the initial image acquisition. Cell viability and killing kinetics were analyzed using Incucyte software, cell analyzer. Images were acquired on SX5- Incucyte instrument (Sartorius AG).

### Expansion of human T Cells for *in vivo* study

T cell isolation from PBMCs. Frozen peripheral blood mononuclear cells (PBMCs) from healthy donors were thawed into ImmunoCult-XF T Cell Expansion Medium (STEMCELL Technologies, Cat# 10981) and immediately washed with MACS Buffer consisting of 0.5% bovine serum albumin (BSA) (Miltenyi Biotec, Cat# 130-091-376) and 2 mM EDTA (Gibco). T cells were isolated by negative selection using the Pan T Cell Isolation Kit (Miltenyi Biotec, Cat# 130-096-535) according to the manufacturer’s instructions. Briefly, cells were incubated with the provided antibody cocktail followed by magnetic microbeads and applied to a magnetic separation column; the unlabeled T cell fraction was collected and washed before culture.Purified T cells were resuspended in ImmunoCult-XF T Cell Expansion Medium (STEMCELL Technologies, Cat# 10981) and activated with the Human ImmunoCult CD3/CD28/CD2 T Cell Activator (STEMCELL Technologies, Cat# 10990) in the presence of interleukin-2 (IL-2) at 50 IU/mL. Cultures were expanded by medium upscaling while maintaining IL-2 at 50 IU/mL: on day 3, culture volume was increased eightfold with fresh medium containing IL-2; on day 5, culture volume was increased fourfold with fresh IL-2. On day 7, cells were harvested and adjusted to 5.0 × 10^7 cells/mL. Expanded T cells were administered in PBS at a dose of 1.0 × 10^7 in cells per mouse via injection on day 7.

### Ex vivo assessment of T-cell functionality in treated mice

Resected tumors were placed in ice-cold medium. Tissues were minced into ∼<5 mm pieces and transferred into GentleMACS C Tubes (Miltenyi Biotec cat# 130-093-237)) containing Tumor Dissociation Kit reagents (Reagents D,R,A) (Miltenyi Biotec, cat#:130-096-730). Samples were mounted on the gentleMACS Dissociator with Heaters (Miltenyi Biotec, catalog 2260) and processed using the recommended program(37C_m_TDK2) for 42 minutes at 37°C. Cell suspensions were passed through 70 µm strainers, rinsed with complete medium (RPMI-1640 + 10% FBS), and centrifuged at 400 × g for 5 minutes at 4°C. Pellets were resuspended in ACK lysis buffer (Gibco, catalog#A1049201) and incubated for 3 minutes at room temperature, quenched with excess medium, and centrifuged as above. Cells were resuspended in 37% Percoll (Cytiva, catalog #17089101) and centrifuged at 400 × g for 20 minutes at room temperature. The resulting cell pellets were collected, washed with MACS buffer (PBS + 1% FBS + 2 mM EDTA), counted, and used for T-cell isolation. Spleens were pressed gently through 70 µm strainers using frosted microscope slides into complete medium. Cell suspensions were centrifuged (400 × g, 5 minutes, 4°C), resuspended in ACK lysis buffer for 3 minutes at room temperature, quenched with medium, and washed once with MACS buffer. Cells were counted and used for T-cell isolation using Miltenyi Biotec human CD2 MicroBeads (cat # 130-091-114) according to the manufacturer’s protocol. Isolated T cells were immediately used for cytotoxicity assays or cytokine profiling.

PC9 cells were seeded in 96-well plates in complete medium. Isolated T cells were added at 1:1 effector:target (E:T) ratios (in the presence or absence of TCE (160 pM) and cultured for 72 hours at 37°C, 5% CO_2_. At endpoint, the plates were centrifuged and adherent PC9 cells were detached with TrypLE (20 minutes at 37°C), neutralized with medium. Cells were stained for viability and relevant markers, and counting beads were added prior to acquisition on a flow cytometer. Tumor cell survival and T-cell activation were quantified by gating on human EpCAM and CD3^+^ T cells.

### Cytokine profiling by intracellular staining

Isolated T cells from tumors or spleens were seeded at 2 × 10^5^ cells per well in 96-well plates in complete medium and stimulated with a T-cell activation cocktail containing Brefeldin A (Biogend, Cat# 423304) for 4 hours at 37°C. Cells were washed with MACS buffer and PBS and stained with a fixable Live/Dead dye for 15 minutes at room temperature in the dark. After washing, Fc receptors were blocked with anti-mouse and anti-human CD16/32 (10 minutes, 4°C). Surface staining was performed for 20 minutes at 4°C. Cells were washed twice with MACS buffer, fixed and permeabilized using eBioscience Foxp3 / Transcription Factor Staining Buffer Set (ThermoFisher, Cat# 00-5523-00). Intracellular cytokine staining (IFN-γ, TNF-α, IL-2) was performed for 30 minutes at room temperature, followed by two washes with Perm/Wash and one wash with MACS buffer. Samples were acquired on a BD FACSymphony A5SE and data were analyzed with FlowJo (v. 10.8.1). Cytokine-positive T-cell frequencies were determined using fluorescence minus one (FMO) control.

### T cell immunophenotyping

Human tumor xenografts were excised and processed using the Tumor Dissociation Kit, mouse (Miltenyi Biotec, cat. no. 130-096-730), in combination with the gentleMACS Dissociator (Miltenyi Biotec, cat. no. 130-093-235), following the manufacturer’s instructions. Dead cells were excluded using a fixable viability dye (Zombie UV, BioLegend) per the manufacturer’s protocol. Fc receptors were blocked with Human TruStain FcXtrade (BioLegend) prior to antibody incubation. Surface staining was performed at 4°C in the dark for 30 minutes using BioLegend Cell Staining Buffer and the following: mouse CD45 (clone 30-F11, BUV395; 1:100), human CD45 (clone HI30, APC-R700; 1:100), CD3 (clone UCHT1, BUV805; 1:100), CD4 (clone SK3, BV750; 1:100), CD8 (clone RPA-T8, BV421; 1:100), CCR7 (clone G043H7, APC-Cy7; 1:100), CD69 (clone FN50, BUV563; 1:100), CD103 (clone Ber-ACT8, RB744; 1:100), CD45RA (clone 5H9, BUV737; 1:100), CD45RO (clone UCHL1, PE-CF594; 1:100), CD7 (clone CD7-6B7, PE-Cy7; 1:100), HLA-DR (clone L243, BV605; 1:100), CD28 (clone CD28.2, BV480; 1:100), CD95 (clone DX2, BUV615; 1:100), CD57 (clone HNK-1, RB545; 1:100), CD39 (clone TU66, BV650; 1:100), PD-1 (clone EH12, BB515; 1:100), Tim-3 (clone F38-2E2, BV711; 1:75), TCF1 (clone C63D9, PE; 1:100), and TOX (clone REA473, APC; 1:100). Human TruStain FcXtrade and Zombie UV were used at manufacturer-recommended dilutions. After surface staining, cells were washed twice with BioLegend Cell Staining Buffer. Staining for transcription factors TCF1 and TOX was performed using the BioLegend Cyto-Fast Fix/Perm Buffer Set (cat. no. 426803) according to the manufacturer’s instructions. Following surface staining and washes, cells were fixed with Cyto-Fast Fix Buffer and permeabilized with Cyto-Fast Perm Buffer, then incubated with anti-TCF1 (C63D9, PE; 1:100) and anti-TOX (REA473, APC; 1:100) for 30 minutes on ice in the dark. Cells were washed and resuspended in BioLegend Cell Staining Buffer for acquisition. Samples were acquired on a Cytek Aurora I-5L spectral flow cytometer equipped with UV (355 nm), violet (405 nm), blue (488 nm), yellow-green (561 nm), and red (640 nm). Data were analyzed in FlowJo v10.10.0 (BD). For dimensionality reduction, tSNE was run on arcsinh-transformed marker intensities using default parameters (perplexity 30, theta 0.5, iterations =1000). Clustering was performed with FlowSOM in FlowJo on the same transformed features, using a 10×10 self-organizing map with hierarchical metaclustering.

### Granzyme B uptake assay

PC9 cells were seeded at in 96-well flat-bottom plates (200 μL) and incubated overnight (37°C, 5% CO2). Cells were treated with ADC 24 h, briefly centrifuged (300 × g, 1 min), and then incubated with inactive Granzyme B (1 μg/mL, 1 h, 37°C). After washing with cold MACS buffer, cells were pelleted (500 × g, 5 min). For flow cytometry, cells were stained with LIVE/DEAD Fixable Aqua (Thermo Fisher L34966; 1:200, 15 min, RT), stained for surface M6PR (APC, BioLegend 364206; 1:100), fixed and permeabilized using eBioscience Foxp3 / Transcription Factor Staining Buffer Set (ThermoFisher, Cat# 00-5523-00), 30 min, 4°C), and stained intracellularly for Granzyme B (BioLegend 372206; FITC1:100, 30 min, 4°C). Samples were washed and acquired. Controls included unstained, single-color, FMO (GrzB), and no-ADC/no-GrzB conditions.

### ATG5 knockdown by siRNA in PC9 cells

PC9 cells were transfected using Lipofectamine RNAiMAX (Thermo Fisher, cat. 13778150) in Opti-MEM I (Gibco, cat. 31985-070). Silencer pre-designed ATG5 siRNA (Thermo Fisher, cat. 4392420; ID s18158; 5 nmol) or Silencer Negative Control siRNA (Thermo Fisher, cat. 4390843) was reconstituted to 100 μM and diluted to 10 μM working stocks. For 96-well plates, siRNA in Opti-MEM was combined 1:1 with RNAiMAX in Opti-MEM to yield 50 μL complexes per well (incubated 5 min at room temperature), targeting a final dose of 1 pmol siRNA and 0.3 μL RNAiMAX per well after cells were added. Complexes were dispensed into wells and overlaid with 50 μL of PC9 cells in antibiotic-free complete medium to a final volume of 100 μL per well. Cells were incubated at 37°C with 5% CO2, and ATG5 knockdown was assessed 24 h post-transfection by qRT-PCR.

### TNFR1 (TNFRSF1A) knockout in PC9 cells

PC9 cells were edited by CRISPR/Cas9 RNP nucleofection. TrueCut Cas9 Protein v2 (Thermo Fisher, cat. A36497) was complexed with TrueGuide synthetic sgRNA targeting TNFRSF1A (Thermo Fisher, cat. A35533; Assay ID CRISPR752924_SGM) for 15 min at room temperature. The RNPs were delivered to 2 × 10^5 PC9 cells using the Lonza 4D-Nucleofector. Cells were immediately transferred to pre-warmed complete RPMI and cultured at 37°C with 5% CO2. Edited cells were cultured for 3 days post-nucleofection, and loss of TNFR1 protein was confirmed by flow cytometry

### Quantitative real-time PCR

Total RNA was extracted using the Zymo Quick-RNA Miniprep Kit (Zymo Research, cat. #R1054) according to the manufacturer’s instructions. On-column DNase digestion was performed, and complementary DNA (cDNA) was synthesized using the High-Capacity cDNA Reverse Transcription Kit (Applied Biosystems, cat. #4374966). Quantitative real-time PCR (qRT-PCR) was performed with TaqMan Fast Advanced Master Mix (Applied Biosystems, cat. #4444557) in 96-well plates. Reactions were run and analyzed on a QuantStudio™ real-time PCR system (Thermo Fisher Scientific). TaqMan Gene Expression Assays (FAM reporter) were used for the following targets: CD120A/TNFR1 (Thermo Fisher, assay ID Hs01042313_m1), CD120B/TNFR2 (Thermo Fisher, assay ID Hs00961750_m1), GAPDH (Thermo Fisher, assay ID Hs02786624_g1), ACTB (Thermo Fisher, assay ID Hs01060665_g1), M6PR (Thermo Fisher, assay ID Hs00974474_m1), and ATG5 (Thermo Fisher, assay ID Hs00355494_m1).

### TCE generation

The TED2 (T-cell Engager DuetMab 2) format is a trivalent bispecific T-cell engager engineered to redirect cytotoxic T cells toward tumor cells. In this format, the molecule typically contains one antigen-binding domain specific for CD3 on T cells and two antigen-binding domains recognizing a tumor-associated antigen (TAA), creating a 2:1 TAA:CD3 binding configuration. Multi-specific modalities were engineered and optimized to have better stability, pairing and developability (41,42).

### MART1 T-Cell Generation

Antigen-specific T cell generation protocol was adapted from(43). HLA-A*02+ PBMCs from cryopreserved naïve cells and fresh peripheral blood were cultured in 6-well G-Rex plates. Dendritic cell differentiation was initiated with IL-4 and GM-CSF on day 1, followed by antigen exposure on day 2 using MART-1 peptide, TNF-α, TLR8 ligand, IL-7, and PGE2. IL-7, IL-15, and IL-2 were subsequently added on day 3 to promote T cell expansion. Following a 3-day expansion phase, on day 6, media supplemented with IL-7, IL-15, and IL-2 was exchanged every 48 hours until day 10. CD8+ T cells were negatively selected via magnetic separation and cryopreserved.

### Generation of M6PR-KO cell line

Guide RNAs (gRNAs) targeting the *M6PR* gene were designed using the CRISPick tool (https://portals.broadinstitute.org/gppx/crispick/public) (44,45) and CHOPCHOP (https://chopchop.cbu.uib.no/) (46). The selected sgRNA sequences were 5′- UUACAGCUUUGAGAGCACUG – 3’ and 5′- GGUGCAAAUCAACAAAAGUA – 3’. Lipofectamine CRISPRMAX (Thermo Fisher Scientific) was used to deliver CRISPR/Cas9 ribonucleoprotein (RNP) complexes into PC9 cells according to the manufacturer’s protocol. After recovery, cells were sorted using an anti-M6PR polyclonal antibody (Invitrogen, Thermo Fisher Scientific, PA5-92876). When single-cell–derived clones reached confluency in a 6-well plate, genomic DNA was extracted using the Puregene Cell Kit (Qiagen). Amplification of the target sequences was conducted using the HotStar Taq Plus Master Mix Kit (Qiagen) per the manufacturer’s protocol. Primers used were 5’ – CACTCAGCCATGACCTCAAA – 3’ and 5’ – TCTCATCCCCCAGTTTTAGC – 3’.

### Statistical analysis

Statistical analyses were conducted in GraphPad Prism 8.3.1. A two-sided unpaired Student’s t-test was used for comparison of two unpaired groups. One-way analysis of variance (ANOVA) with correction for multiple comparisons was used for single comparisons with >2 groups. Two-way ANOVA was used for the analysis of repeated measurements within the groups.

---

**Table 1.** Reagents.

| **Reagent** | **Vendor** | **Catalog number** |
| --- | --- | --- |
| AIM-V | Gibco | 12055-091 |
| Human Serum | GeminiBio | 100-512 |
| Tall G-Rex Plate | Wilson Wolf | 80240M |
| hIL-4 | Stemcell | 78147.1 |
| hGM-CSF | Stemcell | 78140.1 |
| TLR8L | Invivogen | tlrl-lrna40 |
| TNF-a | R&D Systems | 210-TA |
| PGE2 | Stemcell | 72192 |
| IL-2 | R&D Systems | BT002 |
| IL-7 | R&D Systems | BT007 |
| IL-15 | R&D Systems | BT015 |
| EasySep™ Human CD8+ T Cell Isolation Kit | Stemcell | 17953 |
| PBS, pH 7.4 (500 mL) | Sigma-Aldrich | P4474-500ML |
| Anti-Anti (5 mL) | Gibco | 15240-062 |
| RPMI with HEPES modification (450 mL) | Gibco | A10491-01 |
| Penicillin/streptomycin (5 mL) | Gibco | 15140-122 |
| Normocin (1 mL) | InvivoGen | Ant-nr-1 |
| FBS, heat inactivated (50 mL) | Gibco | A56708-01 |
| Low Melting Point Agarose | Invitrogen | 16520100 |
| BOND Dewax Solution | Leica Biosystems | AR9222 |
| BOND Epitope Retrieval ER2 Solution | Leica Biosystems | AR9640 |
| Refine Detection Kit Peroxide Block* | Leica Biosystems | DS9800 |
| Primary Antibody | Santa Cruz Biotechnology | sc-8436 |
| Refine Detection Kit Post Primary* | Leica Biosystems | DS9800 |
| Refine Detection Kit Polymer* | Leica Biosystems | DS9800 |
| Refine Detection Kit Mixed DAB Refine* | Leica Biosystems | DS9800 |
| Refine Detection Kit Mixed DAB Refine* | Leica Biosystems | DS9800 |
| Refine Detection Kit Hematoxylin* | Leica Biosystems | DS9800 |
| TNF-R2 antibody | Cell Signaling Technology | E8D7P |
| IGF-II Receptor/CI-M6PR antibody | Cell Signaling Technology | D8Z3J |

**Table 2.** Flow cytometry antibody and reagents.

| Name | Clone | Fluorochrome | Company | Catalog number |
| --- | --- | --- | --- | --- |
| CD3 | SK7 | APC | BioLegend | 981012 |
| CD3 | OKT3 | PE | BioLegend | 317308 |
| CD4 | SK3 | FITC | BioLegend | 980802 |
| CD4 | L200 | BUV395 | BD Horizon | 564107 |
| CD4 | OKT4 | APC | BioLegend | 317416 |
| CD4 | RPA-T4 | PerCP/Cy5.5 | BioLegend | 300530 |
| CD4 | RPA-T4 | BV785 | BioLegend | 300554 |
| CD4-M | GK1.5 | BV785 | BioLegend | 100453 |
| CD8 | RPA-T8 | BV711 | BD Horizon | 563677 |
| CD8 | RPA-T8 | BV421 | BioLegend | 301036 |
| CD8 | SK1 | FITC | BioLegend | 980908 |
| CD25 | BC95 | BV711 | BioLegend | 302636 |
| CD25 | BC96 | BV510 | BioLegend | 302640 |
| CD25 | BC96 | PerCP | BioLegend | 356132 |
| CD25 | BC96 | PE/Cy7 | BioLegend | 302612 |
| CD44 | BJ18 | PB | BioLegend | 338824 |
| CD45 | 2D1 | FITC | BioLegend | 368508 |
| CD45 | HI30 | PE | BioLegend | 982322 |
| CD45 | HI30 | BV785 | BioLegend | 304048 |
| CD45 | HI30 | BUV395 | BD | 563792 |
| CD45-M | 30-F-11 | PE/Cy7 | BioLegend | 103114 |
| CD69 | FN50 | BV421 | BioLegend | 310930 |
| CD69 | FN50 | APC/Cy7 | BioLegend | 310914 |
| CD69 | FN50 | PE | BioLegend | 302612 |
| CD69 | FN50 | PE | BioLegend | 985202 |
| EPCAM | 9C4 | BV421 | BioLegend | 569568 |
| EPCAM | W18060B | FITC | BioLegend | 312504 |
| TNF-α | MAb11 | APC/CY7 | BioLegend | 502944 |
| IL-2 | MQ1-17H12 | APC | BioLegend | 500310 |
| IFN-gamma | W19227a | PE | BioLegend | 383304 |
| CD120a/TNFR1 | W15099A | APC | BioLegend | 369906 |
| CD120B/TNFR2 | 3G7A02 | PE | BioLegend | 358404 |
| M6PR | QA19A18 | APC | BioLegend | 364206 |
| TER119-M | TER119 | BV650 | BioLegend | 116235 |
| CD31-M | 390 | BV605 | BioLegend | 102427 |
| LC3A/B | D3U4C | PE | Cell signaling | 13611S |
| 7-AAD | NA | NA | BioLegend | 420404 |
| HER2 (CD340) | 24D2 | PE | BioLegend | 324406 |
| TROP2 | 162-46.2 | PE | BioLegend | 363804 |
| EGFR | AY13 | PE | BioLegend | 352904 |
| Granzyme B | QA16A02 | FITC | BioLegend | 372206 |
| Granzyme B | GB11 | BV421 | BioLegend | 396414 |
| PD-1 (CD279) | EH12.2H7 | APC/Cy7 | BioLegend | 367416 |
| PD-L1 (CD274) | 29E.2A3 | BV421 | BioLegend | 329714 |
| Mouse IgG2a $\kappa$ isotype control | MOPC-173 | APC | BioLegend | 400222 |
| CD120b/TNFR2 | FAB226 | PE | BioLegend | 358404 |
| Rat IgG2a $\kappa$ isotype control | RTK2758 | PE | BioLegend | 400508 |
| LIVE/DEAD Fixable Blue | NA | DAPI channel | Thermo Fisher | L34962 |
| LIVE/DEAD Fixable Aqua | NA | BV480-510 | Thermo Fisher | L34966 |
| LIVE/DEAD Fixable Near-IR | NA | APC/Cy7 | Thermo Fisher | L34994 |
| Rat IgG1 isotype control | RTK2071 | BV421 | BioLegend | 400430 |

**Table 3.** Other antibodies and reagents.

| Name | Clone | Company | Catalog number |
| --- | --- | --- | --- |
| Recombinant anti-CD3 | OKT3 | BioLegend | 317347 |
| Recombinant anti-CD28 | CD28.2 | BioLegend | 302943 |
| Recombinant IL-2 | NA | Sigma-Aldrich/Roche | 11147528001 |
| Human TNF-α blocking mAb | 1825 | R&D Systems | MAB210-100 |
| Anti-human TNF-α (Infliximab biosimilar) | Infliximab | Bio X Cell/VWR | SIM0006 |
| Human TRAIL/TNFSF10 blocking mAb | 75411 | R&D Systems | MAB375-500 |
| Human NKG2D/CD314 blocking mAb | 149810 | R&D Systems | MAB139-100 |
| Human Fas Ligand/TNFSF6 blocking mAb | NOK-1 | R&D Systems | AB126 |
| Anti-HMGB1 (Ultra-LEAF) | 3E8 | BioLegend | 651414 |
| IFN-γ blocking mAb | mAb7 (InvivoGen) | InvivoGen | hifng-mab7-02 |
| IFNAR1 blocking mAb | Anifrolumab | AZ | AZ |
| Purified Rat anti-mcd16/cd32 | 2.4g2 | Biolegend | 553142 |
| UltraComp eBeads | NA | invitrogen | 01-3333-42 |
| ArC-reactive beads | NA | invitrogen | A10346A |
| TNF-α Human ELISA Kit | NA | Thermo Fisher | KHC3011 |
| IFN-γ Human ELISA Kit | NA | Thermo Fisher | KHC4021 |

