## Supplemental Information for "Mechanisms regulating combination effect of antibody-drug conjugates and cancer immunotherapy"

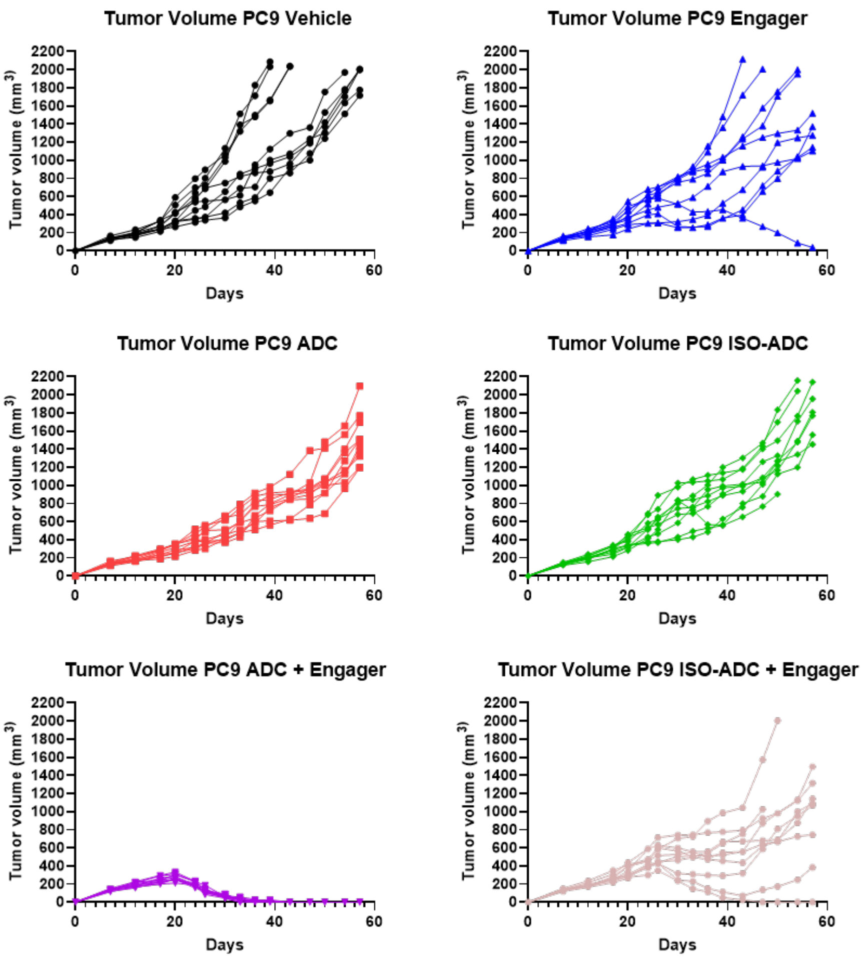


**Figure S1. Antitumor effect of ADC/TCE combination therapy in vivo.** NSG mice bearing PC9 tumors were treated with ADC, TCE and their combination as described in Figure 1H. Individual mouse tumor growth curves are shown.

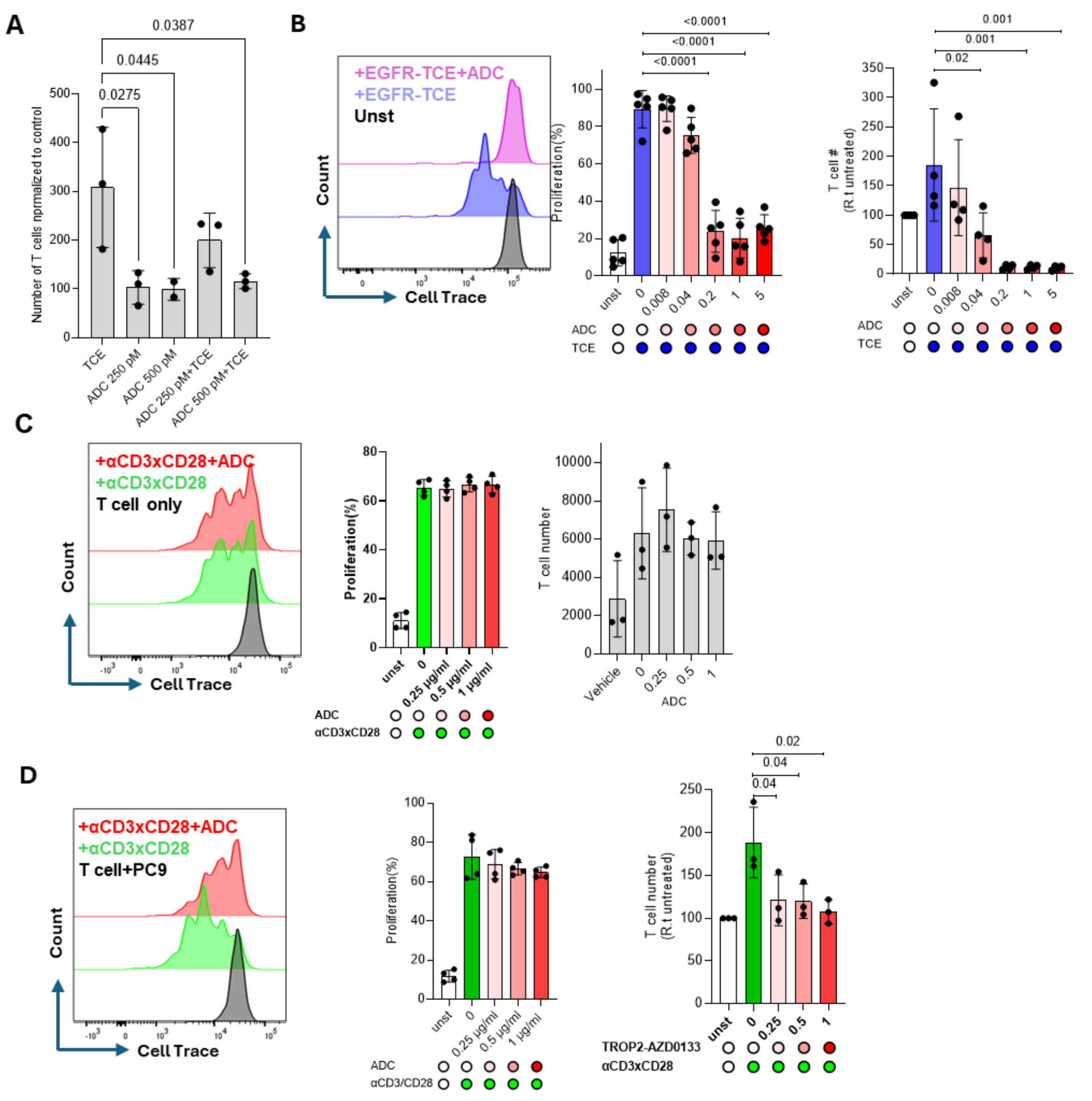


**Figure S2. Toxic effect of ADC on T cells. A**. T-cell numbers in cultures PC9 tumor cells and T cells in the presence of ADC payload, TCE, or their combination. Counts were normalized to untreated controls (considered as 100) (n=3). **B.** T cells proliferation and number in the presence of PC9 tumor cells treated with TCE (160 pM) in combination with indicated ADC concentrations. Left panel - representative flow-cytometry plots of T-cell proliferation. Middle panel - percentage proliferation (n=5). Right panel – number of T-cells (n=4). The number of T cells were normalized to untreated control. **C.** T-cell proliferation in the absence of cancer cells. T cells were stimulated with ImmunoCult™ Human CD3/CD28 25 µl/ml in the presence of different concentrations of ADC. Left panel - representative flow-cytometry plot. Middle panel - percentage proliferation (n=4) Right panel: T-cell number (n = 3). **D.** T-cell proliferation and number in the presence of cancer cells. T cells were co-cultured with PC9 cells and stimulated with CD3/CD28 (1 µg/ml) in the presence of different concentrations of ADC. Left panel - representative flow-cytometry plots. Middle panel: percentage of proliferating T cells (n=4). Right panel – number T cells normalized to untreated control (n = 3). Individual values, mean, and s.d. are shown. P values were calculated one-way ANOVA test with correction for multiple comparisons.

**A**

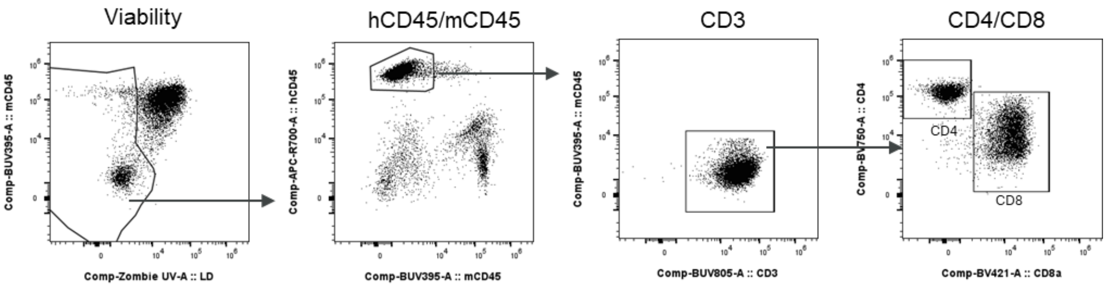


**B**

| Tumor | Spleen |
|---|---|
| CD28 | CD28 |
| HLA-DR | HLA-DR |
| CD45RA | CD45RA |
| TIM3 | TIM3 |
| PD1 | PD1 |
| TCF1 | TCF1 |
| CD103 | CD103 |
| CD7 | CD7 |
| CD69 | CD69 |
| TOX | CD39 |
| | CD45RO |
| | CD103 |
| | CD7 |
| | CD69 |
| | CCR7 |
| | CD95 |
| | TOX |

**Figure S3. Expression of surface molecules on spleen and tumor T cells after the treatment of mice with ADC.** Mice were inoculated with tumor cells and 5 days later ADC and T cells were administered. The TCE was administered 7 days after the first T-cell injection. Two days after the second injection, tumors and spleens were collected for T-cell characterization. **A.** Representative flow-cytometry plots illustrating the gating strategy used to define T-cell subsets. **B.** Markers that were not changed in T cells between TCE and TCE + ADC groups.

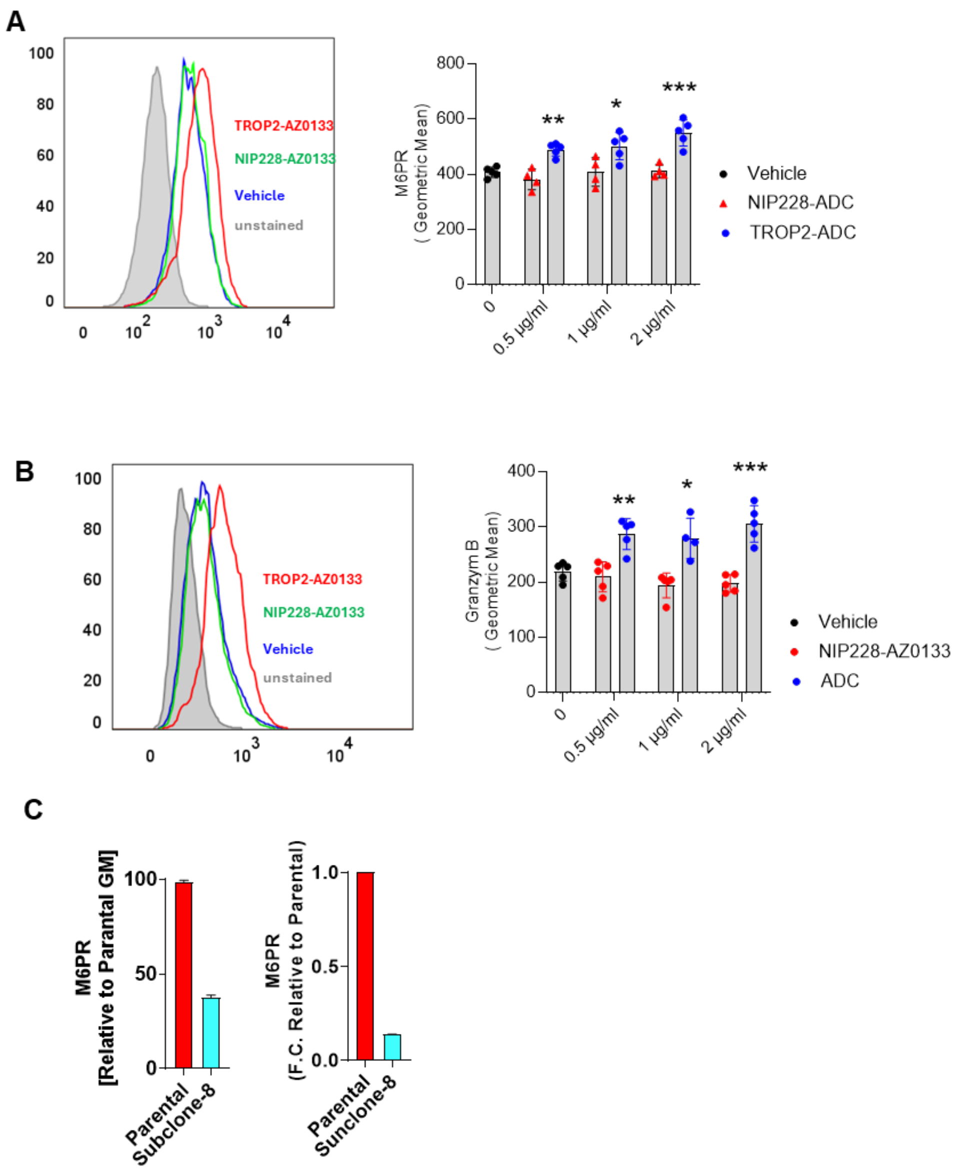


**Figure S4. M6PR is not involved in enhanced efficacy of the ADC-TCE combination. A**. M6PR surface expression on cancer cells after treatment with ADC or control NIP228-AZ0133. Left panel - representative flow-cytometry histograms. Right panel - geometric mean fluorescence intensity (gMFI) for M6PR (n=4). **B.** Uptake of recombinant granzyme B by cancer cells cultured alone, with ADC, or NIP228-AZ0133. Left panel - representative flow-cytometry plots. Right panel - gMFI of intracellular granzyme B staining. (n=5). **C.** M6PR knockout in PC9 cells. Left panel surface M6PR expression by flow cytometry in PC9-M6PR-KO vs parental cells, Right panel - M6PR gene expression relative to parental PC9 cells. P values were calculated using unpaired, two-sided Student's t-tests. * $p < 0.05$, ** $p < 0.01$; *** $p < 0.001$

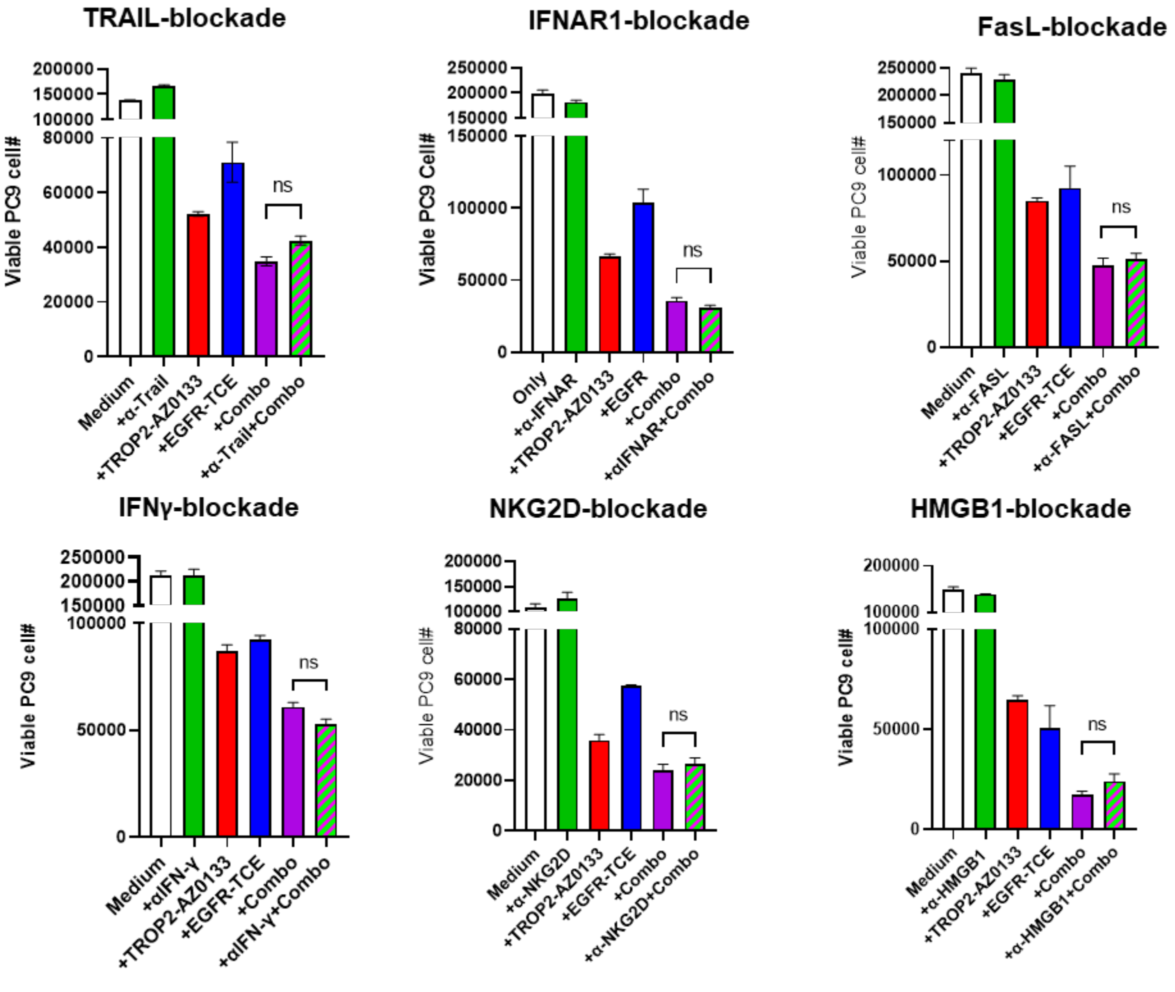


**Figure S5. Effect of neutralization of various T-cell derived cytokines on antitumor activity of ADC-TCE combination.** Viable PC9 tumor cell counts after co-culture with T cells in the presence of TCE (160 pM), ADC (0.04 µg/ml), or their combination, with indicated antibodies. Cultures were performed at a 1:1 T-cell : target ratio for 3 days. Antibodies were present throughout the co-culture period. Concentration 1.25 µg/ml was used after titration of the highest effective concentration based on prior reports. In all graphs n=3, mean and SD are shown. P values were calculated using one-way ANOVA test with corrections for multiple comparisons.

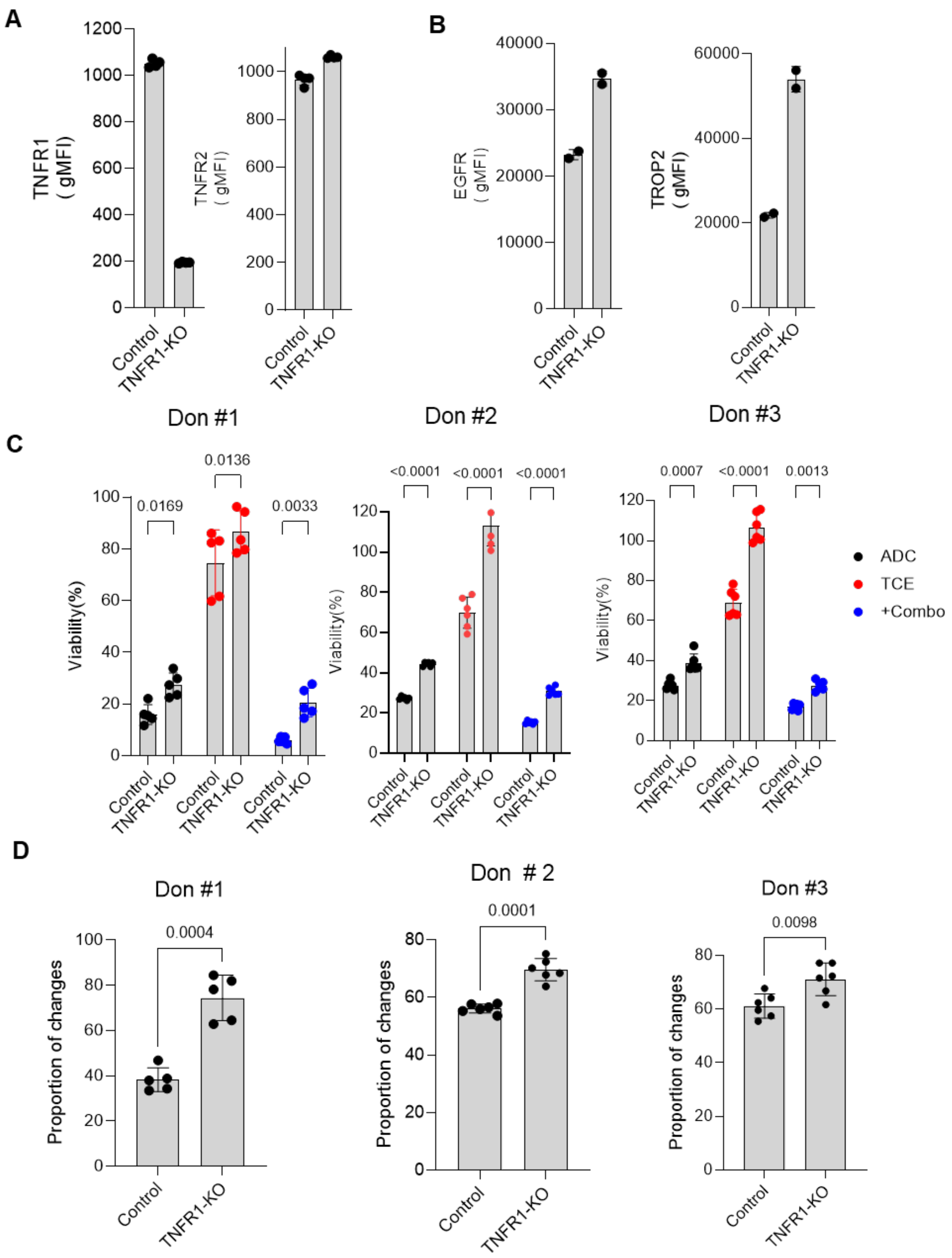


**Figure S6. Effect of TNFR1 deletion on ADC-TCE antitumor activity. A.** Surface expression of TNFR1 (left) and TNFR2 (right) on PC9 and PC9-TNFR1-KO cells, measured by flow cytometry (n = 3). **B.** Surface expression of target molecules—TROP2 (left) and EGFR (right) in PC9 and PC9-TNFR1-KO cells, measured by flow cytometry (n=2) **C.** Viability of PC9 and PC9-TNFR1-KO cells co-cultured with T cells in the presence of TCE (160 pM), ADC (0.04

μg/ml), or the ADC+TCE combination. Data are shown for individual donors (n=5). **D**. Ratio of tumor-cell killing in PC9 and PC9-TNFR1-KO cells treated with the ADC+TCE combination, calculated by normalizing to the untreated control within each corresponding group (n=6). Individual values, mean, and s.d. are shown. P values were calculated using unpaired, two-sided Student's t-tests.

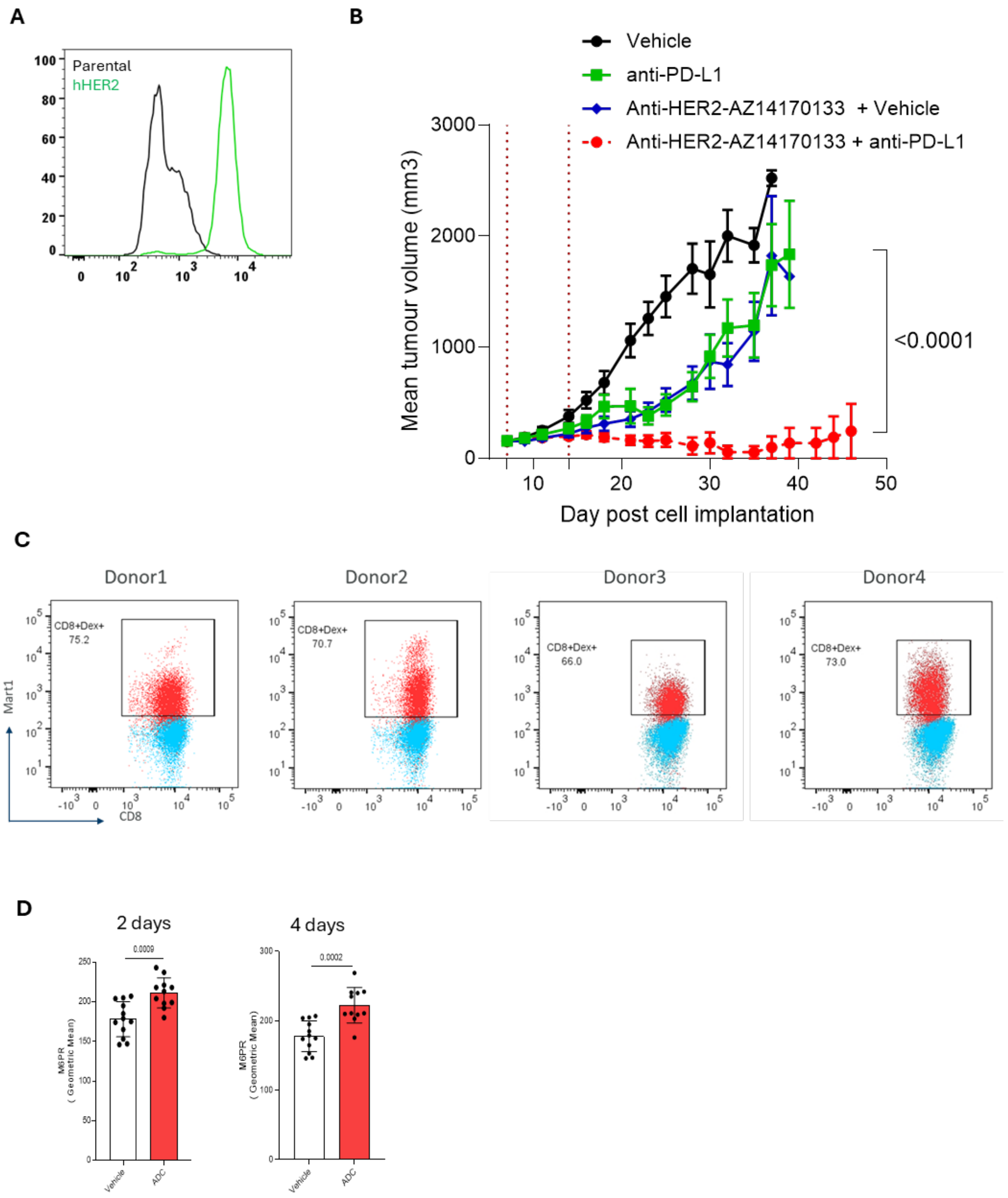


**Figure S7. Antitumor activity of ADC-TCE combination in mice bearing EMT6 tumor expressing HER2. A.** Expression of h-HER2 EMT6 tumor cells after transfection. **B.** Treatment of EMT6-huHER2 tumor bearing mice with HER2-AZ14170133 ADC and PD-L1. EMT6-huHER2 cells were implanted into syngeneic mice C57BL/6 mice. Animals were treated with HER2-AZ14170133 on days 7 and 14, and with anti–PD-L1 on days 7, 10, 14, and 17. Tumor volumes were measured at regular intervals, and animals were euthanized when tumor burden exceeded ethical limits. N=9. P values were calculated in two-way ANOVA test. **C.** MART1 dexamer staining of T cells generated by repeated stimulation with peptide loaded DCs. Results with cells from individual donors are shown. **D.** Expression of M6PR on tumor cells in vivo following ADC treatment as described in **Fig. 4C**. Panels depict analyses at 2 and 4 days after the first ADC dose. Each point represents individual mouse. N=9. Individual results, mean, and SD are shown. P values were calculated in two-sided Student's t-tests.

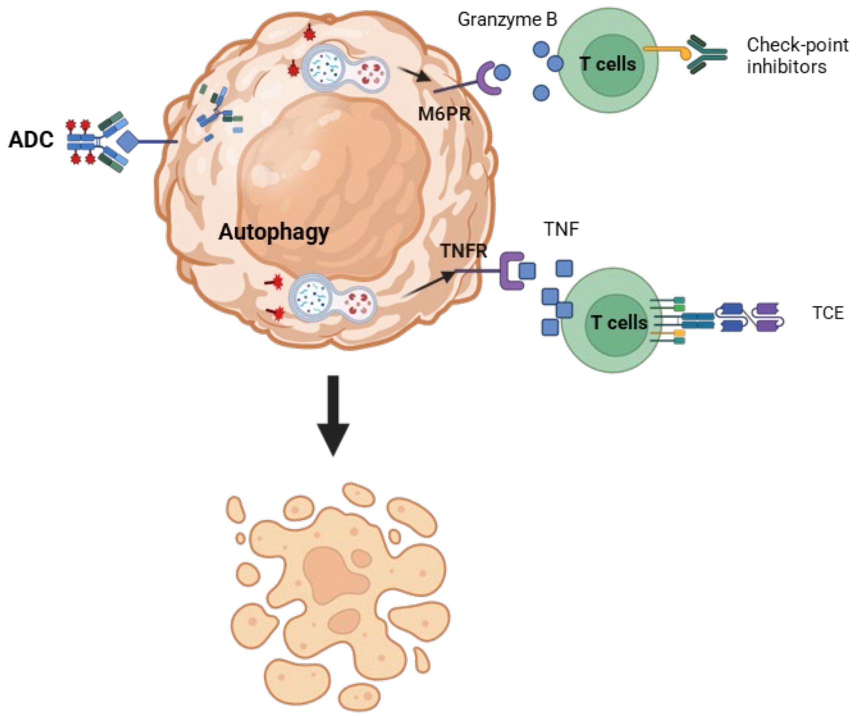


**Figure S8. Schema of the mechanisms of combination effect of ADC and immunotherapy of cancer.** Payload released from ADC induces autophagy in tumor cells leading to up-regulation (via redistribution) of TNFR and M6PR on the cell surface. This makes tumor cells more susceptible to killing by activated T cells depending on the nature T cell activation. TCE stimulated T cells produce large amount of TNFα that kill tumor cells via TNFR bindings, whereas antigen-specific T cells mediate increased killing via release of granzyme B that binds to M6PR.
